# Stable coexistence in plant-pollinator-herbivore communities requires balanced mutualistic vs antagonistic interactions

**DOI:** 10.1101/2021.06.29.450358

**Authors:** Youssef Yacine, Nicolas Loeuille

## Abstract

Ecological communities consist of multiple species interacting in diverse ways. Understanding the mechanisms supporting coexistence requires accounting for such a diversity. Because most works focus either on mutualism or predation, how pollination and herbivory interactively determine the stable coexistence in plant-pollinator-herbivore communities is still poorly understood. Studying the typical three-species module of such communities, we determine the conditions allowing stable coexistence then investigate how its maintenance constrains the relative interaction strengths. Our results show that coexistence is possible if pollination is sufficiently strong relative to herbivory, while its stability is possible if herbivory is sufficiently strong relative to pollination. A balance between pollination and herbivory is therefore required. Interestingly, shared preferences for plant phenotypes, that would favor such balance, have been frequently reported in the empirical literature. The identified ecological trade-off between attracting pollinators and deterring herbivores therefore also appears as an emergent property of stable plant-pollinator-herbivore communities.

## 1. Introduction

Multiple species interacting closely together form an ecological community. A topic of long-standing interest in community ecology is to understand what mechanisms drive the coexistence of species and its maintenance over time. It is now well established that the way ecological interactions connect species - the type of interactions, the network topology as well as the distribution of interaction strengths - plays a decisive role. Combining modelling approaches with empirical data, several works indicate for instance that weak trophic interactions are crucial to maintain the stability of complex food webs (McCann et al., 1998; Neutel et al., 2002). Deriving general laws is, however, difficult. The network properties and topologies favoring the maintenance of coexistence indeed vary with the type of interaction characterizing the community, mutualism or antagonism in particular (Thébault and Fontaine, 2010). The ecological processes and structural patterns supporting the maintenance of coexistence within single-interaction-type communities can, moreover, considerably differ from the ones at play within communities with several interaction kinds (e.g. Mougi and Kondoh, 2012; Sauve et al., 2014). Studies of such communities should therefore significantly improve our understanding of ecological communities, especially given that most species get simultaneously involved in a diversity of interaction networks (Fontaine et al., 2011; Kéfi et al., 2012). Most terrestrial plant species (≈ 90% of flowering plants, Ollerton et al., 2011), for instance, are involved in a mutualistic interaction with their animal pollinators, while suffering from herbivorous predation (antagonism). Plant-pollinator-herbivore communities are, in addition, of particular interest due to their critical role in agricultural production (Klein et al., 2007; Oerke, 2006), as well as the serious threats global change poses to them (Atwood et al., 2020; Potts et al., 2010). The study of stable coexistence within these communities is thus of high applied relevance while offering the opportunity to gain new conceptual insights into the functioning of mutualistic-antagonistic communities.

Understanding stable coexistence within plant-pollinator-herbivore communities requires explicitly accounting for both the mutualistic (i.e. plant-pollinator) and the antagonistic (i.e. plant-herbivore) interaction. A large body of empirical evidence indeed documents non-additive effects of pollination and herbivory on plant densities, in both uncultivated (Gómez, 2005; Herrera, 2000; Herrera et al., 2002; Pohl et al., 2006) and cultivated (Lundin et al., 2013; Strauss and Murch, 2004; Sutter and Albrecht, 2016) plant species. The strength of the mutualistic interaction is affected by the antagonistic interaction and vice versa, explaining such an interactive effect. Herbivores may, for instance, preferentially consume plant species bearing abundant flowers or developing fruits as a result of strong pollination (Herrera, 2000; Herrera et al., 2002). By decreasing floral display, herbivore damages can reduce pollination (Adler et al., 2001; Cardel and Koptur, 2010; Pohl et al., 2006). In addition to floral display, herbivory-induced changes in plant chemistry can also deter pollinators (Kessler et al., 2011).

Indirect interactions between two species within a community can also be mediated by their effect on the density of a third species (Wootton, 2002). Ubiquitous in natural communities, such indirect effects play a key part in the maintenance of coexistence (Burns et al., 2014; Menge, 1995). By isolating the structural building blocks of complex ecological networks - modules or motifs - it becomes easier to unravel such indirect effects and their implications for community maintenance (Milo et al., 2002). Modules are therefore small sets of interacting species characteristic of the studied community, whose study enables deeper insights into the mechanisms at play at the broader scale (Milo et al., 2002; Stouffer and Bascompte, 2010).

In plant-pollinator-herbivore communities, the typical module consists of two animal species - a pollinator and a herbivore - sharing a common resource plant species (Fig. 1A.a). The resulting indirect interaction between pollinators and herbivores is an antagonism (Fig. 1A.a, Holland *et al.* 2013). Pollinators allow the community to sustain a higher herbivore density by increasing plant productivity while herbivores, on the other hand, decrease pollinator density by reducing resource availability. Theoretical works indicate that the presence of pollinators can even make the herbivore population viable (Georgelin and Loeuille, 2014; Mougi and Kondoh, 2014a). As illustrated by Georgelin & Loeuille (2014), direct vs. indirect ecological effects can be of similar magnitude. Their study indeed reports a constant herbivore density despite increasing herbivore mortality. In their model, over a wide range of herbivore mortalities, the direct mortality-induced losses on herbivores are totally offset by the indirect gain resulting from a higher pollinator density consecutive to the herbivorous predation release on plants. Further increasing mortality, however, leads to the abrupt collapse of the herbivore population, which illustrates that combining different interactions also has important implications in terms of community stability (Mougi and Kondoh, 2014b).

**Figure 1:**
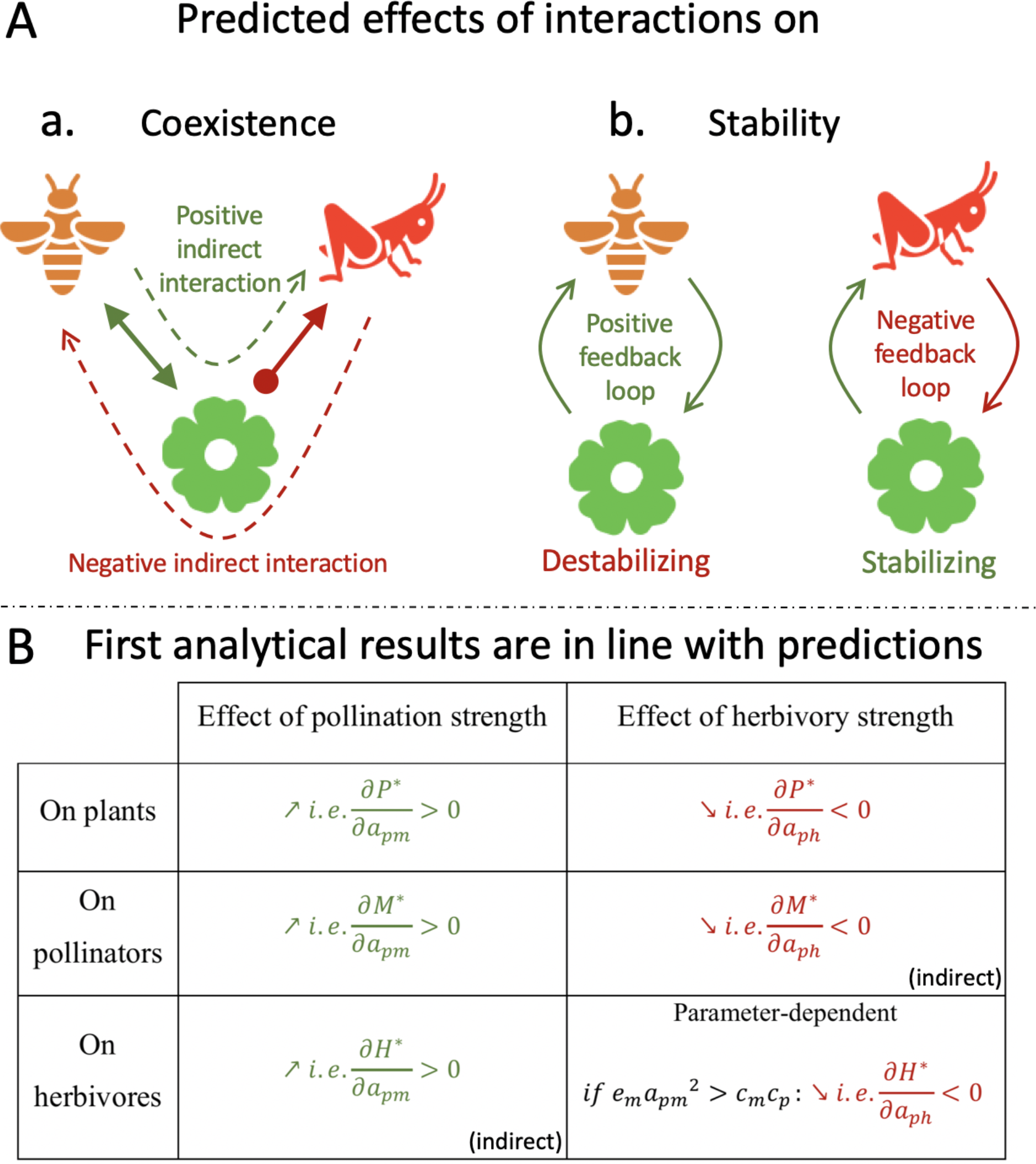
**A. Predicted effects of interactions on stable coexistence. a.** Predicted effects on densities. Solid (resp. dashed) arrows for direct (resp. indirect) interactions. **b.** Predicted effects on stability. **B. Variation of biomass densities at stable coexistence with the strength of interactions.** These first analytical results are in line with predictions (see A.a). Analytical proofs in appendix B.IV.

Mutualisms, such as pollination, intrinsically entail positive feedback loops (Fig. 1A.b). Positive feedbacks are destabilizing (Levins, 1974; Neutel and Thorne, 2014) as they tend to amplify the direct effect of a perturbation. As a result, unstable behaviors have been identified in theoretical models of mutualism, including tipping points (Lever et al., 2014) or unbounded population growths driven by an “orgy of mutual benefaction” (May, 1981). The latter is, however, seldom observed in nature. One possible explanation is that antagonistic interactions, such as predation, could prevent this behavior in real systems. Negative feedback loops born from antagonistic interactions (Neutel and Thorne, 2014) could restore stability by counterbalancing the positive loops arising from mutualisms (Fig. 1A.b). This hypothesis implies that the relative magnitude of pollination vs. herbivory plays a critical role, which is in line with the findings of several theoretical investigations on mutualistic-antagonistic modules (Georgelin and Loeuille, 2014; Holland et al., 2013; Mougi and Kondoh, 2014b; Sauve et al., 2016a).

The goal of the present paper is to understand how stable coexistence within plant-pollinator-herbivore communities constrains the relative strengths of pollination and herbivory, i.e. the relative per capita effects of each interacting animal species on plant population growth. In contrast with most previous theoretical works on mutualistic-antagonistic modules, the relationships governing stable coexistence we obtain are analytical. Such relationships between pollination and herbivory are derived from the population dynamics of the characteristic three-species module (Fig. 1A.a), in which both animal intake rates (i.e. functional responses) are assumed linear to achieve analytical tractability. We discuss such an assumption at the end of the present work (section 4). Finally, the per-capita effect of plant-animal interactions on community dynamics is mediated by animal densities, which in turn depend on other ecological parameters such as animal mortalities or intraspecific competition rates. We therefore extend our analysis by studying their influence, which confirms the robustness of our results. In what follows, we show that stable coexistence within plant-pollinator-herbivore communities requires a balance between the strengths of pollination and herbivory. Such a pattern ensues from the opposite effect each interaction has on coexistence and stability (Fig. 1A). Coexistence is favored by pollination and disfavored by herbivory, as a result of both direct and indirect ecological interactions (Fig. 1A.a). Stability is enhanced by herbivory and reduced by pollination, as a result of the respective feedback loops (Fig. 1A.b). It is interesting to note that a large body of empirical literature (e.g. Irwin *et al.* 2003) reports shared preferences for plant phenotypes between pollinators and herbivores that would favor balanced interactions, which appear here as an emergent property of stable plant-pollinator-herbivore communities.

## 2. Model presentation

### 2.1 Ecological dynamics

We formulate the dynamics of the biomass densities of three interacting species - a plant P, a pollinator M, and a herbivore H - using ordinary differential equations:

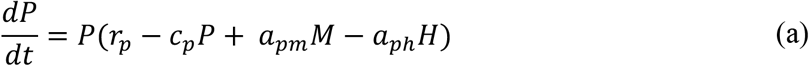

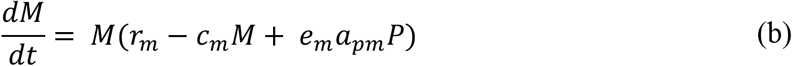

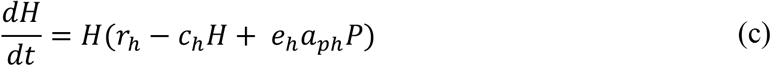

Plants have a positive intrinsic growth rate (*r*_*p*_ > 0, autotrophs), while both pollinators (*r*_*m*_ < 0) and herbivores (*r*_*h*_ < 0) have a negative one (heterotrophs). As in previous models (e.g. Sauve *et al.* 2014), we thus assume the plant-animal interaction to be obligate for animals and facultative for the plant. Intraspecific competition is accounted for. The animal competition rates (*c*_*m*_, *c*_*h*_) correspond to interference while, for the plant species (*c*_*p*_), this rate essentially captures the competition for resources such as light, water, and nutrients (Craine and Dybzinski, 2013). Interspecific interactions, whose strength is *a*_*pm*_ for pollination and *a*_*ph*_ for herbivory, affect population growths proportionally to biomass densities. The use of a linear functional response for mutualism exposes the model to unbounded population growths (May, 1981). It, however, enables testing whether this behavior could be top-down controlled by herbivory, placing our work in the line of research tackling how the community context could explain the stability of mutualisms in nature (e.g. Ringel *et al.* 1996). Finally, *e*_*m*_ and *e*_*h*_ are the conversion efficiencies from plants to animals. Parameter details are given in table 1.

**Table 1:** List of all model parameters and variables with their biological significance, value, and dimension (***M*** for mass, ***L*** for length, and ***t*** for time).

| Variables and parameters |  | Biological meaning | Value | Dimension |
| --- | --- | --- | --- | --- |
| *Variables* | $P$ | Plant biomass density | | $M \cdot L^{-2}$ |
| | $M$ | Pollinator biomass density | | $M \cdot L^{-2}$ |
| | $H$ | Herbivore biomass density | | $M \cdot L^{-2}$ |
| *Interaction strength* | $a_{pm}$ | Strength of pollination<br>(i.e. per capita effect of pollinators on plant population growth) | [0,3] | $(M \cdot L^{-2})^{-1} \cdot t^{-1}$ |
| | $a_{ph}$ | Strength of herbivory<br>(i.e. per capita effect of herbivores on plant population growth) | [0,3] | $(M \cdot L^{-2})^{-1} \cdot t^{-1}$ |
| *Other ecological parameters* | $r_p$ | Plant intrinsic growth rate | 10 | $t^{-1}$ |
| | $r_m$ | Pollinator intrinsic growth rate | [-5, -1] | $t^{-1}$ |
| | $r_h$ | Herbivore intrinsic growth rate | [-5, -1] | $t^{-1}$ |
| | $c_p$ | Plant intra-specific competition rate | 0.6 | $(M \cdot L^{-2})^{-1} \cdot t^{-1}$ |
| | $c_m$ | Pollinator intra-specific competition rate | [0.2,0.6] | $(M \cdot L^{-2})^{-1} \cdot t^{-1}$ |
| | $c_h$ | Herbivore intra-specific competition rate | [0.2,0.6] | $(M \cdot L^{-2})^{-1} \cdot t^{-1}$ |
| | $e_m$ | Plant to pollinator conversion efficiency | [0.1,0.3] | Dimensionless |
| | $e_h$ | Plant to herbivore conversion efficiency | [0.1,0.3] | Dimensionless |

### 2.2 Ecological equilibria

When the three population growth rates vanish (equations (a b c) are null), we reach an ecological equilibrium (*P*\*, *M*\*, *H*\*). At equilibrium, each population can either be present or absent which leads to 8 potential equilibria (expressions in Appendix B.III). The present work focuses on the equilibrium in which the three species are present, hereafter “the coexistence equilibrium”. We study under which conditions of interaction strengths - (*a*_*pm*_, *a*_*ph*_) - this equilibrium corresponds to positive biomass densities (i.e. is feasible) and stable (i.e. is resilient to small perturbations). See Appendix B.I for detailed definitions.

### 2.3 Two-population subcommunities

The plant-pollinator-herbivore community is constituted of two subcommunities – plant-pollinator and plant-herbivore – sharing the same plant species. Such subcommunities have extensively been studied in the literature (e.g. Goh 1976; Vandermeer & Boucher 1978). We briefly report here their dynamics (see Appendix B.II for details).

The plant-herbivore subcommunity is characterized by one feasible and globally stable equilibrium. Either the plant population at carrying capacity 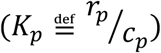 can support the herbivore population and both coexist, or the herbivores go extinct while plants persist.

Two distinct regimes are possible for the plant-pollinator subcommunity, depending on the comparison between pollination strength and self-limiting competitions (Appendix B.II.2). (1) If the pollination positive feedback loop (destabilizing, Fig. 1A.b) is smaller than that from competition, there is one feasible and globally stable equilibrium. This equilibrium corresponds to plant-pollinator coexistence when the carrying capacity of plants is large enough to support the pollinator population. Otherwise, plants persist while pollinators go extinct. (2) If the pollination positive feedback loop is stronger than that from competition, unbounded population densities are possible. In this case, when the carrying capacity of plants is sufficient to make pollinators viable, populations unboundedly grow irrespective of initial densities. Otherwise, unbounded growth is observed if initial densities are large enough while only plants persist if it is not the case.

## 3. Results

At the coexistence equilibrium when feasible and stable, all biomass densities increase with the strength of pollination (Fig. 1B). On the contrary, both plant and pollinator densities decrease as herbivory gets stronger, while herbivore density can either increase or decrease (Fig. 1B). Matching our predictions (Fig. 1A.a), these dynamics are illustrated in Fig. 2, which shows how densities depend on herbivory for three pollination levels (Fig. 2a-b-c), and on pollination for three herbivory levels (Fig. 2d-e-f). Fig. 2 especially demonstrates that population dynamics are determined by both pollination and herbivory interactively. For instance, the decline of herbivore density with the strength of herbivory is observed when pollination is strong (Fig. 1B & 2c), which we interpret as a consequence of the strong indirect antagonism with pollinators (Fig. 1A.a). Another example is that the strength of one interaction affects the level the other interaction has to reach in order for the focal animal to persist in the community. As pollination increases, the minimal level of herbivory allowing herbivores to persist gets lower (Fig. 2a vs 2b). Pollination favors the feasibility of coexistence. On the contrary, herbivory disfavors the feasibility of coexistence. As herbivory gets stronger, the minimal level of pollination allowing pollinators to persist gets higher (Fig. 2d&e vs 2f).

**Figure 2:**
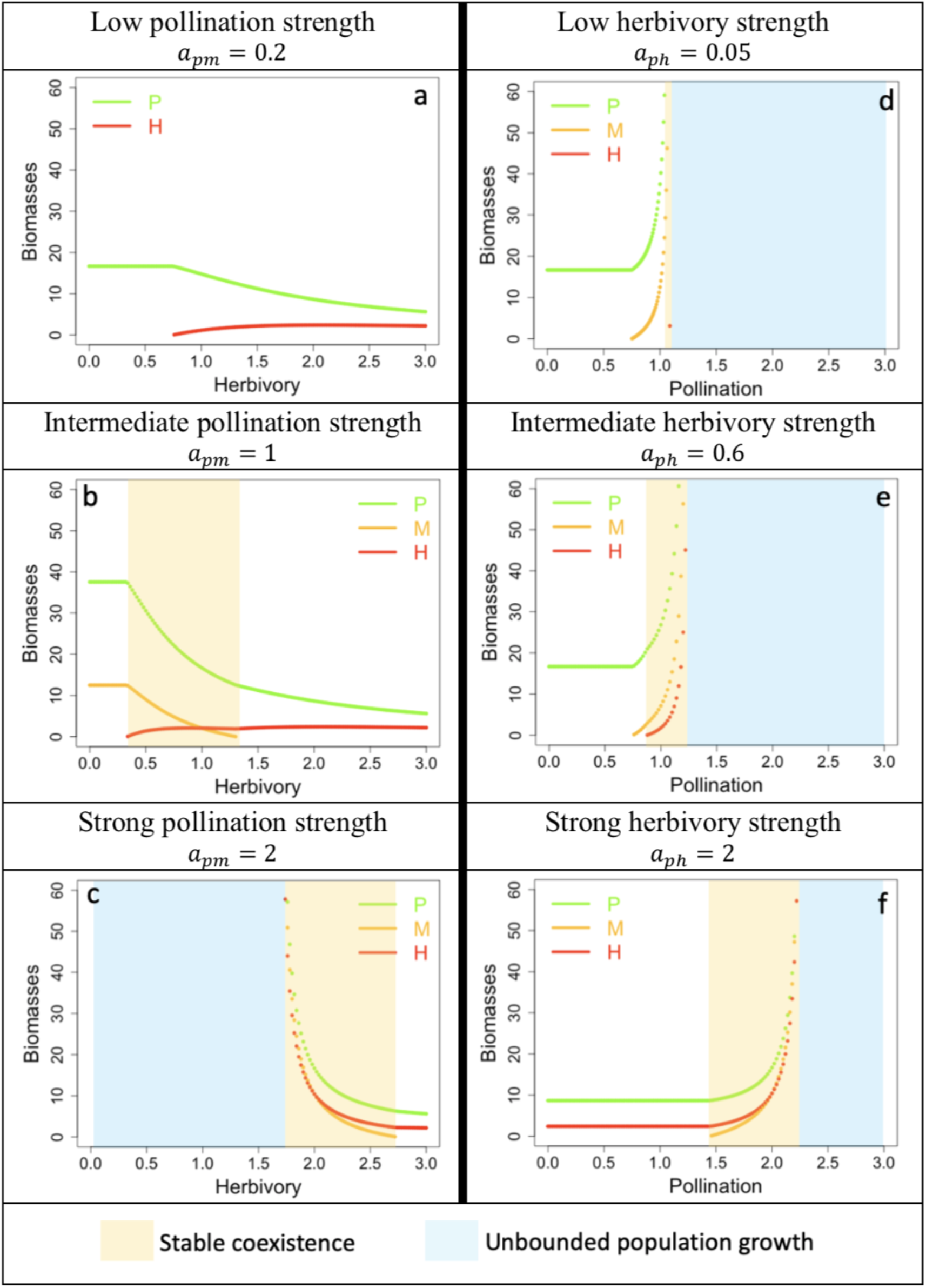
Biomass densities at equilibrium depend interactively on pollination and herbivory strengths. The curves are determined analytically (see appendix B.III). **a-b-c** Dependence of densities on herbivory for three pollination levels. In **a**, pollination intensity is too low for pollinators to persist. **d-e-f** Dependence of densities on pollination for three herbivory levels. Parameter set: *r*_*p*_ = 10, *r*_*m*_ = *r*_*h*_ = −2.5, *c*_*p*_ = 0.6, *c*_*m*_ = *c*_*h*_ = 0.4, *e*_*m*_ = *e*_*h*_ = 0.2.

In line with our predictions (Fig. 1A.b), stability displays opposite patterns: it is favored by herbivory and disfavored by pollination. For a given level of herbivory, populations display unstable dynamics for high pollination strengths (Fig. 2 d-e-f, blue background). This instability captures the unbounded growth of biomass densities driven by the mutualism. As herbivory gets stronger, higher pollination levels are needed for unbounded growth to happen (Fig. 2d vs 2e vs 2f). Herbivory can indeed restore stability (Fig. 2c): starting from an initially unbounded situation, increasing herbivory restores finite densities.

The strength of pollination contributes positively to the feasibility of coexistence and negatively to its stability. It is the opposite for herbivory. Although presented for a given parameter set (Fig. 2), these two main results are general as they derive from the analytical relationships governing stable coexistence (table 2). They imply that stable coexistence requires a balance between the strengths of pollination and herbivory to achieve both feasibility and stability. Such a balance can be observed in Fig. 2: as one interaction gets stronger, the range of the other interaction intensities allowing stable coexistence shifts toward larger values (orange background, Fig. 2b vs 2c, Fig. 2e vs 2f).

**Table 2:** Analytical relationships governing stable coexistence. The fourth column indicates how each relationship is affected by the strength of interactions (favored +, disfavored −). Note that the third and fourth columns present a simplified summary of our analysis (see subsequent text and Appendix C, especially tables S3 & S4). Notations: 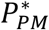 plant density at plant-pollinator equilibrium; 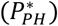 plant density at plant-herbivore Nc NO equilibrium; “*num”* numerator; “*den*” denominator. Finally, the interplay between pollination and herbivory is difficult to disentangle in relationship (2), which led us to distinguish two cases (inequality (2’a) for (a) and (2’b) for (b) below). An increase in pollination (*a*_*pm*_) makes the relationship shift from (a) to (b). In (b), 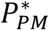 loses its biological significance as the plant-pollinator subcommunity grows unboundedly. 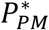 is, in this case, a mathematical function (as defined in Appendix C.II.2), which explains its counterintuitive behavior with the variation of pollination strength *.

| Relationship | Mathematical meaning | Biological interpretation | Effect of interaction strengths |
| --- | --- | --- | --- |
| (1)<br>$r_m + e_m a_{pm} P_{PH}^* \geq 0$ | Feasibility<br>(num($M^*$) $\geq 0$) | Pollinators can invade the plant-herbivore community | $a_{pm} +$ (direct)<br>$a_{ph} - (P_{PH}^* \searrow)$ |
| (2)<br>$(c_p c_m - e_m a_{pm}^2)(r_h + e_h a_{ph} P_{PM}^*) \geq 0$ | Feasibility<br>(num($H^*$) $\geq 0$) | (a) Stable plant-pollinator dynamics with herbivores able to invade the plant-pollinator community<br><u>or</u><br>(b) Plant-pollinator orgy with bounded herbivory | (a) $a_{pm} + (P_{PM}^* \nearrow)$<br>$a_{ph} +$ (direct)<br>(b) $a_{pm} + (P_{PM}^* \searrow)^*$<br>$a_{ph} -$ (direct) |
| (3)<br>$c_h e_m a_{pm}^2 - c_p c_m c_h - c_m e_h a_{ph}^2 < 0$ | Feasibility<br>(den($M^*, H^*$) $\geq 0$)<br>Stability | Total feedback at level 3 is negative | $a_{pm} -$<br>$a_{ph} +$<br>(feedback loops, Fig. 1b) |
| (4)<br>$(c_p P^* + c_m M^* + c_h H^*)(P^* M^* (c_p c_m - e_m a_{pm}^2) + P^* H^* (c_p c_h + e_h a_{ph}^2) + M^* H^* c_m c_h) - P^* M^* H^* (c_h c_m c_p - c_h e_m a_{pm}^2 + c_m e_h a_{ph}^2) > 0$ | Stability | Negative feedback at level 3 is weaker than the product of negative feedback at lower levels (1 & 2) | Undetermined |

### 3.1 Relationships governing stable coexistence

Positive animal densities necessarily imply a positive plant density because animals are obligate plant-interactors (Appendix C.I.1). In other words, the coexistence equilibrium is feasible if, and only if, both animal species have positive densities, which leads to two inequalities. It is stable if, and only if, all three eigenvalues of the Jacobian matrix (Appendix B.III.1) calculated at the coexistence equilibrium have a negative real part, which is equivalent to the three Routh-Hurwitz inequalities (Appendix B.III.3). One of these inequalities is satisfied if feasibility is assumed. Therefore, there are four relationships (i.e. inequalities) that are necessary and sufficient for the stable coexistence of plants, pollinators and herbivores (Appendix C.II). These relationships, as well as their biological interpretations, are presented in Table 2. We illustrate the biological implications underlying them using Fig. 3, which indicates the community composition depending on the strengths of pollination and herbivory.

**Fig. 3:**
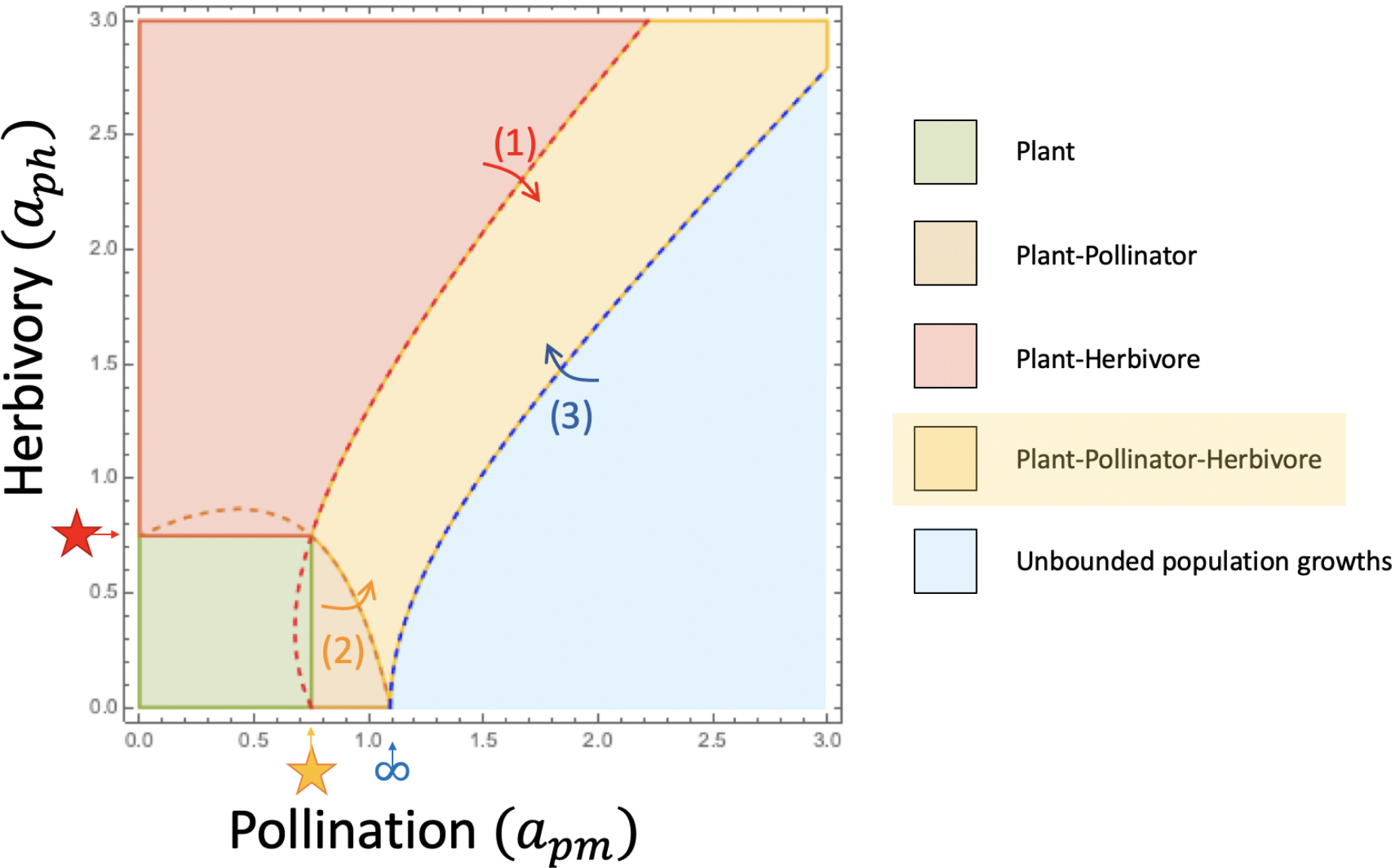
Stable coexistence requires balanced pollination vs. herbivory strengths. Given the strength of pollination (*a*_*pm*_) and herbivory (*a*_*ph*_), the stable equilibria are determined and the point of the graph is colored accordingly. In blue, no equilibrium is stable so densities grow unboundedly. Arrows (1), (2), and (3) indicate the transitions enabling the satisfaction of relationships (1), (2), and (3) (**Table 2**), indicated by the dashed red, orange, and blue curve, respectively. These three relationships are sufficient to achieve stable coexistence given the parameter set (as in **Fig. 2**), indicating that relationship (4) is less constraining here. The orange (resp. red) star indicates the level of pollination (resp. herbivory) that makes pollinators (resp. herbivores) viable when only plants are present (hence at carrying capacity). Unbounded growth is possible in the plant-pollinator subcommunity when the strength of pollination is higher than the level figured by an infinity symbol. Note that stable coexistence (yellow area) requires the two interactions to be of similar magnitude.

Assuming stability (relationship (3) actually), coexistence is feasible if and only if relationships (1) and (2) are satisfied.

Relationship (1) indicates that the per capita growth rate of pollinators, when low in density and within a plant-herbivore community at ecological equilibrium, is positive (Appendix C.II.1). Pollinators are thus able to invade the plant-herbivore community so that this relationship governs the transition (red dotted curve, arrow (1)) between the plant-herbivore equilibrium (red) and the coexistence equilibrium (yellow) in Fig. 3. Besides, the plant density within the plant-herbivore community 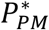 decreases when herbivory (*a*_*ph*_) intensifies (Appendix C.II.1). In such a situation, pollination (*a*_*pm*_) has to get stronger as well in order for pollinators to invade the plant-herbivore community (relationship (1’)).

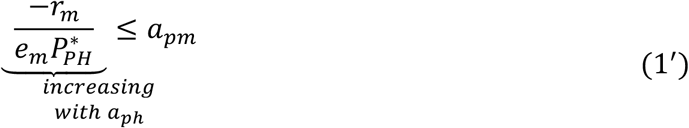

The interpretation of the second relationship depends on whether unbounded population growth is possible or not within the plant-pollinator community, i.e. on the competition loop (*c*_*p*_*c*_*m*_) being weaker or stronger than the pollination feedback loop 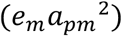.

If unbounded growth is not possible, the relationship indicates that the per capita growth rate of herbivores, when low in density and within a plant-pollinator community at ecological equilibrium, is positive. In this case, the relationship governs the transition (orange dotted curve, arrow (2)) between the plant-pollinator equilibrium (orange) and the coexistence equilibrium (yellow) in Fig. 3. We mathematically demonstrate that in such a case, the feasibility of coexistence implies its global stability (Appendix B.IV). Relationships (1) and (2) are thus necessary and sufficient for stable coexistence (Fig. 3, left side of ∞). Furthermore, stronger pollination (*a*_*pm*_) makes herbivores viable at lower predation intensities (*a*_*ph*_) (relationship (2’a)) due to a higher plant density within the plant-pollinator community 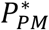 (Appendix C.II.2).

If unbounded growth is possible, the relationship sets an upper limit to the strength of herbivory (relationship (2’b)), which we interpret as a condition for herbivores to not exclude pollinators by reducing plant biomass too strongly. In fact, relationship (2’b) (feasibility of *H*\*) is critical for a parameter configuration over which the persistence of herbivores is due to the presence of pollinators (Appendix C.II.2, Fig. S3). In such parameter instances, 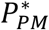 loses ibtisological meaning (as the plant-pollinator equilibrium is unstable) and decreases with pollination (*a*_*pm*_), and alternative stable states are possible (Fig. S3). Note that no transition corresponds to relationship (2’b) in Fig. 3 as relationship (2) is only constraining at the left of the infinity symbol (∞) for the given parameter set.

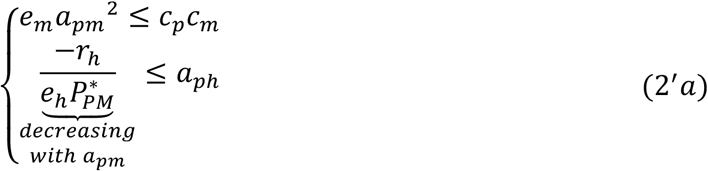

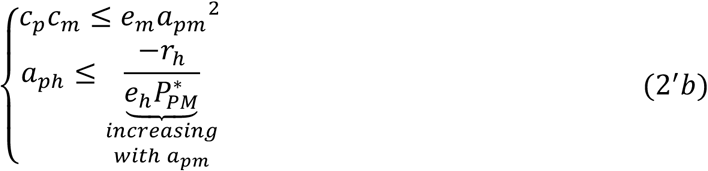

Relationships (1’) and (2’) clearly show that pollination favors the feasibility of coexistence while herbivory disfavors it. Both relationships indeed tend to be satisfied when pollination strengthens or herbivory weakens.

Assuming feasibility, coexistence is stable if and only if relationships (3) and (4) are satisfied.

Relationship (3) corresponds to the total feedback at level 3 (i.e. summation of the strengths of all three-element combinations of non-overlapping feedback loops, details in Appendix C.II.3) being negative. Pollination disfavors stability by contributing positively to this feedback, while it is the opposite for herbivory. Stability requires the competitive and the herbivory feedback loops to overcome the pollination feedback loop. Relationship (3’) emphasizes the consecutive constraint limiting pollination. It governs the transition (blue dotted curve, arrow (3)) from unbounded growth (blue) to stable coexistence (yellow) in Fig. 3.

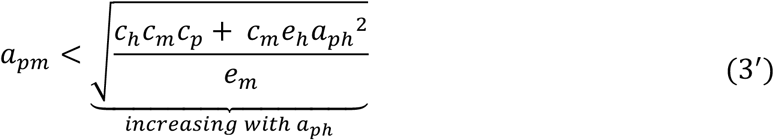

Relationship (4) is harder to interpret. Given that relationships (3) and (4) imply that the feedback at each level is negative, relationship (4) could be interpreted as proposed by Levins (1974): the negative feedback with long time lags (level 3) is weaker than the shorter-loop negative feedback (level 1 & 2) (details in Appendix C.II.4). Also, the constraints imposed by this relationship on interaction strengths are not analytically tractable, due to the effect of interactions on equilibrium densities.

By combining relationships (1’) and (3’), we obtain a necessary condition for stable coexistence (relationship (5)) which implies a positive correlation between pollination and herbivory. Stable coexistence within plant-pollinator-herbivore communities requires a balance between the strengths of pollination and herbivory. Stable coexistence in Fig. 3 (yellow) therefore happens around the first diagonal, where pollination and herbivory are of similar magnitudes.

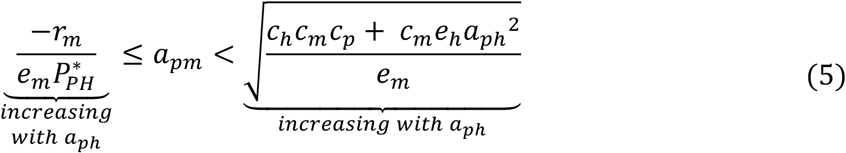

### 3.2 Other ecological parameters also affect stable coexistence

In addition to the per capita effect of plant-animal interactions (i.e. interaction strength), stable coexistence depends on the densities of animal species, which in turn depend on their intrinsic growth and competition rates, as well as their conversion efficiencies. We consequently study the effect of animal growth rates (*r*_*m*_ vs. *r*_*h*_, Fig. 4A & Fig. S4, appendix D), animal competition rates (*c*_*m*_ vs. *c*_*h*_, Fig. 4B & Fig. S5, appendix D) and conversion efficiencies (*e*_*m*_ vs. *e*_*h*_, Fig. S6, appendix D) on community composition. This investigation also constitutes a robustness check as we vary the parameters that were fixed hitherto.

**Fig. 4:**
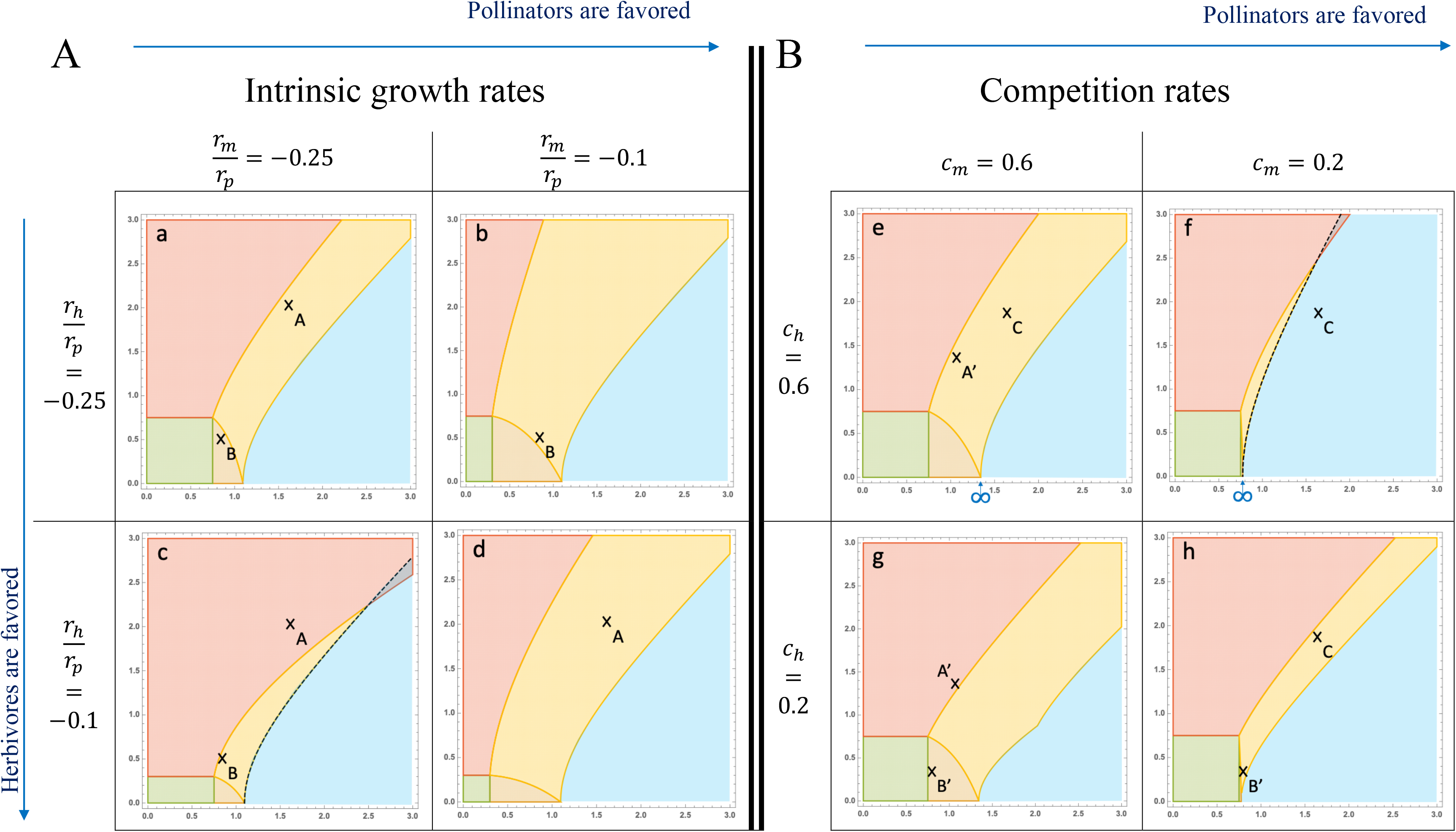
Stable coexistence also depends on animal intrinsic growth rates (A) and competition rates (B). <u>X-axis:</u> Pollination (*a*_*pm*_); <u>Y-axis:</u> Herbivory (*a*_*ph*_) <u>Color legend:</u> 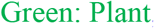, 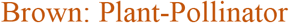 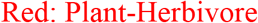, 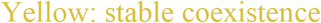 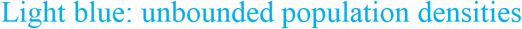. In c & f, alternative states are observed (red and blue overlap): if plant and pollinator densities are initially above a given threshold (dependent on initial herbivore density), populations grow unboundedly; otherwise, pollinators are excluded. Parameters: *r*_*p*_ = 10, *c*_*p*_ = 0.6, *e*_*m*_ = *e*_*h*_ = 0.2; (A) *c*_*m*_ = *c*_*h*_ = 0.4; (B) *r*_*m*_ = *r*_*h*_ = −2.5. Note that Fig. S4 and S5 extend the results of 4A and 4B, respectively.

First of all, when stable coexistence is possible, it happens when the strengths of pollination and herbivory are approximately of the same magnitude (yellow area, Fig. 4 and S4 & S5 & S6 in Appendix D), as analyzed above (relationship (5)).

Stable coexistence is facilitated when the pollinator intrinsic growth rate is higher than the herbivore one. The range of pollination and herbivory strengths allowing stable coexistence indeed gets wider on the upper right of Fig. 4A. The explanation relies on two points: the feasibility of coexistence is favored by pollination and disfavored by herbivory; intrinsic growth rates play a major part in the feasibility of coexistence (relationships (1)&(2)), but only a minor part in its stability (no effect on relationship (3)). Analytical details are available in Appendix D. I. An increase in the pollinator growth rate makes it easier for pollinators to invade the plant-herbivore community (Fig. 4c vs 4d, point A). Due to a higher plant density, herbivores can also invade the plant-pollinator community more easily (Fig. 4a vs 4b, point B). Likewise, a higher herbivore growth rate enables an easier invasion of the plant-pollinator community by herbivores (Fig. 4a vs 4c, point B). It, however, makes the invasion of the plant-herbivore community by pollinators harder due to the reduction of plant density (Fig. 4a vs 4c, point A). Note finally that despite similar growth rates, the community is endangered when these growth rates are too low (Fig. S4, Appendix D).

Stable coexistence is facilitated when competition is stronger among pollinators than among herbivores. The range of pollination and herbivory strengths allowing stable coexistence indeed gets wider in the lower-left of Fig. 4B. Such a pattern is due to the effect of competition rates on stability (relationship (3)), which is much stronger than their effect on feasibility (relationships (1) & (2)). Analytical details are available in Appendix D.II. As competition among herbivores gets stronger, the plant density within the plant-herbivore community increases as a result of predation release. It becomes easier for pollinators to invade (Fig. 4g vs 4e, point A’). Unbounded dynamics are, however, facilitated (Fig. 4h vs 4f, point C) because the positive destabilizing loop increases more than the negative stabilizing loops (relationship (3)). In the plant-pollinator community, a lower pollinator density ensuing from a stronger competition rate is responsible for a lower plant density. It thus becomes harder for herbivores to invade (Fig. 4h vs 4g, point B’). Stability is, however, enhanced due to the stronger control of the pollination positive feedback in both the plant-pollinator subcommunity (Fig. 4f vs 4e, infinity symbol) and the three-species community (Fig. 4f vs 4e, point C).

To summarize, the results obtained from studying the effect of these other parameters support our main results, i.e. pollination favors feasibility at the expense of stability while it is the opposite for herbivory. Indeed, any parameter variation that benefits pollinators (higher growth rate *r*_*m*_, weaker competition *c*_*m*_ or higher conversion efficiency *e*_*m*_ (Appendix D.III)) favors feasibility, disfavors stability or both. Likewise, any parameter variation that benefits herbivores (higher growth rate *r*_*h*_, weaker competition *c*_*h*_ or higher conversion efficiency *e*_*h*_ (Appendix D.III)) disfavors feasibility, favors stability or both.

## 4. Discussion

At the core of community ecology, understanding the mechanisms that support the maintenance of species coexistence is of primary importance in a time of major threats to biodiversity due to global changes (Barnosky et al., 2011). In food webs, it has been shown that the coupling of weak and strong trophic interactions was among such mechanisms (McCann et al., 1998; Neutel et al., 2002). Because weak links can dampen the oscillatory dynamics ensuing from strong links, this unbalanced interaction pattern promotes stable coexistence. In contrast, we suggest that in mutualistic-antagonistic communities, a balance between the strengths of the two interaction types is required to achieve stable coexistence. This main result of our study is in agreement with the findings of several previous theoretical investigations on mutualistic-antagonistic communities, both at the module (Holland et al., 2013; Sauve et al., 2016a) and the network (Mougi and Kondoh, 2012) scale.

The balance between pollination and herbivory is driven by the opposite effects each type of interaction has on coexistence (i.e. feasibility) and stability.

In line with theoretical findings (Georgelin and Loeuille, 2014; Mougi and Kondoh, 2014b; Sauve et al., 2016a), we show that pollination increases herbivore density by enhancing plant density, while the effect of herbivory on pollinators is utterly opposite. This remains true when mutualism is modeled as a modified consumer-resource interaction, thus accounting for exploitative competition between animal species (Holland *et al.* 2013). Congruent direct effects on plant densities have been confirmed by several field experiments (Herrera, 2000; Herrera et al., 2002; Sutter and Albrecht, 2016), but empirical documentation of the consecutive indirect ecological effects between herbivore and pollinator species remains weak (e.g. Gómez 2005).

In contrast with feasibility, we find stability to be favored by herbivory and disfavored by pollination, in line with the theory on feedback loops (relationship (3), Levins 1974). Several studies have indeed shown that pollination networks are prone to display unstable dynamics, such as sudden collapses consecutive to the crossing of tipping points (Dakos and Bascompte, 2014; Kaiser-Bunbury et al., 2010; Lever et al., 2014), as positive feedbacks amplify and propagate disturbances. The important role of predation (herbivorous here) in stabilizing population dynamics, on the other hand, has early been identified (Menge and Sutherland, 1976; Nicholson, 1954; Oksanen et al., 1981). Our results confirm that the consecutive negative feedback can stabilize the dynamics of mutualistic-antagonistic communities. It is important to note, however, that the effects of each interaction type on the stability of such communities are inconsistent across models (Georgelin and Loeuille, 2014; Holland et al., 2013; Sauve et al., 2016a). The different assumptions on the variation of the animal intake rates with plant density (i.e. functional responses) largely explain such contrasting results. It is nonetheless frequent to observe that the stability of the whole community is driven by the subcommunity displaying stable dynamics when considered in isolation. Yet, unstable dynamics are possible when merging two stable subcommunities as shown by Mougi & Kondoh (2014b). In their work, cycling densities are reported, driven by a delayed plant recovery after its exploitation by herbivores. The delay ensues from the fact that most of the productivity gain from pollination is captured by herbivores, which might be particularly problematic in an agricultural context, especially given that it has been reported in empirical studies several times (Gómez, 2005; Herrera et al., 2002). An integrative management of pollination and biological control can, fortunately, enable synergetic interactions between ecosystem services (Sutter and Albrecht, 2016).

It is important to highlight that instability, in our model, encompasses two behaviors whose biological implications are utterly different: (1) the loss of one or several species (Fig. 3, red-brown-green areas) vs. (2) the unbounded growth of population densities (Fig. 3, blue area) driven by an “orgy of mutual benefaction” (May, 1981). While coexistence is not maintained in the first case, it is in the second case. Another notion of stability – permanence (Hutson and Schmitt, 1992) - enables to distinguish between these two cases: a biological community is said to be permanent if the densities of all species are always above a minimal threshold. Unbounded population growth is thus a case of “permanent coexistence” (Hutson and Law, 1985), a concept that captures the diversity of population dynamics that permit the coexistence of species in real biological communities. The orgy of mutual benefaction is, however, seldom observed in nature in spite of mutualisms being widespread (Bronstein, 1994). This indicates that the assumptions of simple models of mutualism are likely violated in real biological systems. The functional response, which we assume linear for both interactions in order to gain analytical tractability, could saturate at high pollination levels when the handling time becomes limiting (e.g. Soberon & Martinez Del Rio 1981). The community context can also impede unrealistic population growth (Freedman et al., 1987; Heithaus et al., 1980; Ringel et al., 1996). While intraspecific competitions prevent this behavior up to a given level of pollination (Holland et al., 2002), we show here that the presence of a third species – the herbivore – allows for even stronger pollination levels to be compatible with biologically relevant finite population densities (relationship (3)). It is thus not surprising that orgies of mutual benefaction are not observed in nature as any two-species mutualism displaying such dynamics would accumulate enemies until restoring the balance required for stable coexistence. Several mechanisms could underlie this community assembly process. Firstly, as the plant biomass is booming, more and more herbivore species are becoming viable in the focal patch (e.g. relationship (2’a)). Because the plant population defines the threshold beyond which herbivore species can invade, as the plant density grows, the filter existing on the possible herbivore community weakens, and more herbivores species are susceptible to come and control the dynamics. Secondly, existing trophic links would likely strengthen as a result of adaptive foraging on the booming plant species in response to its abundance increase relative to other available resources. Adaptive foraging has notably been proposed as an important stabilizing process within complex trophic networks (Kondoh, 2003). In particular, Mougi & Kondoh (2014a) show how the interplay between adaptive foraging, pollination, and herbivory can support the maintenance of stable coexistence in plant-pollinator-herbivore communities.

Empirical evidence suggesting a balance between pollination and herbivory in natural communities does exist. At the module scale, several experimental studies manipulating the presence of animal species find the effects of pollination and herbivory on plant fecundity to be roughly of the same magnitude, approximately canceling each other (Gómez, 2005; Herrera, 2000; Herrera et al., 2002; Sutter and Albrecht, 2016). At the network scale, Melián *et al.* (2009) show that most strong interactions, mutualistic and antagonistic, are concentrated in the same few plant species of the Doñana Biological Reserve (Spain). Sauve *et al.* (2016b) exhibit a positive correlation between the number of pollinators and herbivores that interact with a given plant of the Norwood farm (UK). In line with our results, this correlation contributes positively to the stability of the community. Our results also imply that cascades of extinctions may be expected within plant-pollinator-herbivore networks as a result of the current global pollinator decline (Potts et al., 2010), given the weakening of pollination relative to herbivory.

Empirical evidence linked to species traits also supports the idea of a balanced interaction pattern. Indeed, a large number of studies documents shared preferences for plant phenotypes between pollinators and herbivores. Favoring balanced pollination vs. herbivory, shared preferences have been reported for a large number of plant traits, including flower color (Irwin et al., 2003), floral display (Cariveau et al., 2004; Gómez, 2003), chemical volatiles (Andrews et al., 2007; Theis et al., 2014), nectar quantity (Adler and Bronstein, 2004) or reproductive system (Asikainen and Mutikainen, 2005). Such a pattern implies that plant species are subject to an ecological trade-off between attracting pollinators and deterring herbivores (Strauss et al., 2002, 1999). Our work indicates that this trade-off might be ubiquitous as it fosters the stable coexistence of plant-pollinator-herbivore communities, explaining why it has been reported across a broad diversity of plant taxa. Traits of plant species might be subject to conflicting selection arising from such a trade-off (Strauss and Whittall, 2006), with potentially important implications in terms of diversity maintenance. In the case of the wild radish *Raphanus sativus,* for instance, it has been shown that the maintenance of a flower color dimorphism (white vs. pink) was due to both the pollinators and the herbivores interacting preferentially with white morphs (Irwin et al., 2003; McCall et al., 2013; Stanton, 1987). The question of whether such dimorphism emerged, in the first place, because of the interplay between pollination and herbivory, remains open. The study of mutualistic-antagonistic communities, plant-pollinator-herbivore in particular (Strauss and Irwin, 2004), thus offers opportunities to significantly improve our understanding of the ecological processes supporting the coexistence of species in natural systems, but also of the complex eco-evolutionary dynamics driving the maintenance of biodiversity.

## Acknowledgments

The authors would like to thank Prof. Sharon Y. Strauss, and Dr. François Duchenne, for their helpful feedback on the manuscript.

