## Supporting Information (Appendices) for "Stable coexistence in plant-pollinator-herbivore communities requires balanced mutualistic vs antagonistic interactions"

Appendix A: Nondimensionalization…………………………...2

Appendix B: Study of ecological dynamics…………………....3

I. Preliminary definitions……………………………………….3

II. Subcommunities dynamics ………………………………..3

III. Ecological equilibria in the plant-pollinator-herbivore community………………………………………………………..…4

IV. First analytical results………………………...…………7

Appendix C: Relationships governing stable coexistence…..10

I. Characterization of stable coexistence………………………10

II. Relationships governing stable coexistence………………..11

Appendix D: The effect of the other parameters on stable coexistence…………………………………………………..20

I. Animal intrinsic growth rates………………………………..20

II. Animal intraspecific competition rates……………………..22

III. Animal conversion efficiencies…………………………….24

### Appendix A: Nondimensionalization

In order to improve the readability of the mathematical analyses presented in the supporting information, the model can be rewritten in a nondimensionalized but equivalent way, which allows for fewer parameters. The analyses that use this reformulation are indicated by a star (*) accompanying the title of the corresponding section.

The dynamical system studied follows the equations:

$$\frac{dP}{dt} = P(r_p - c_p P + a_{pm} M - a_{ph} H)$$

$$\frac{dM}{dt} = M(r_m - c_m M + e_m a_{pm} P)$$

$$\frac{dH}{dt} = H(r_h - c_h H + e_h a_{ph} P)$$

It can be rewritten:

$$\frac{dP'}{dt'} = P'(1 - P' + \alpha_{pm} M' - \alpha_{ph} H')$$

$$\frac{dM'}{dt'} = M'(\rho_m - M' + \lambda_m \alpha_{pm} P')$$

$$\frac{dH'}{dt'} = H'(\rho_h - H' + \lambda_h \alpha_{ph} P')$$

With:

$$(P', M', H', t') = (\frac{c_p}{r_p} P, \frac{c_m}{r_p} M, \frac{c_h}{r_p} H, r_p t)$$

$$(\rho_m, \rho_h) = \left(\frac{r_m}{r_p}, \frac{r_h}{r_p}\right) < (0,0)$$

$$(\lambda_m, \lambda_h) = \left(\frac{e_m c_m}{c_p}, \frac{e_h c_h}{c_p}\right)$$

$$(\alpha_{pm}, \alpha_{ph}) = (\frac{a_{pm}}{c_m}, \frac{a_{ph}}{c_h})$$

### Appendix B: Study of ecological dynamics

#### I. Preliminary definitions

##### 1. Feasibility

Our dynamical system (equations (a, b, c)) captures the dynamics of a plant-pollinator-herbivore community as long as the variables, which are biomass densities, are positive. An ecological equilibrium is therefore biologically meaningful if it corresponds to positive densities, i.e. if it is feasible.

##### 2. Stability

Local stability

What we call stability in this work corresponds to local stability. An equilibrium is locally stable when the population dynamics tend to return to the equilibrium following a small perturbation (resilience). Local stability is technically characterized by the fact that all eigenvalues of the Jacobian matrix at the equilibrium have a strictly negative real part, or by the equivalent Routh-Hurwitz criterion (Murray 2002). The latter consists of deriving three coefficients from the Jacobian Matrix ($a_1, a_2, a_3$). The equilibrium is locally stable if and only if: $a_1 > 0$; $a_3 > 0$; $a_1 a_2 - a_3 > 0$.

Global stability

Global stability implies local stability. An equilibrium is globally stable when the population dynamics tend to return to the equilibrium following a perturbation, irrespective of its intensity. Note that perturbations that imply the extinction of one or more populations are excluded.

#### II. Subcommunities dynamics

##### 1. Plant-Herbivore subcommunity

Only one equilibrium is feasible and globally stable, depending on the parameter set. If the plant population, when at carrying capacity (i.e. $\frac{r_p}{c_p}$), is sufficient for the herbivore population to have a positive growth rate when low in density (i.e. $r_h + e_h a_{ph} \left[\frac{r_p}{c_p}\right] > 0$) then stable coexistence between plants and herbivores is observed after a transient state. If not, the herbivore population goes extinct and the plant population remains at carrying capacity.

#### 2. Plant-Pollinator subcommunity

The subcommunity can behave according to two regimes depending on the relative strength between the self-limiting competition loop ($c_p c_m$) and the positive feedback loop due to pollination ($e_m a_{pm}^2$).

If competition overcomes pollination (i.e. $c_p c_m - e_m a_{pm}^2 > 0$), population densities are always bounded. In such a case, there is exactly one feasible and globally stable equilibrium. If the plant population, when at carrying capacity ($\frac{r_p}{c_p}$), is sufficient for the pollinator population to have a positive growth rate when low in density (i.e. $r_m + e_m a_{pm} \left[\frac{r_p}{c_p}\right] > 0$) then stable coexistence between plants and pollinators is observed after a transient state. If not, the pollinator population goes extinct and the plant population remains at carrying capacity.

If not (i.e. $c_p c_m - e_m a_{pm}^2 < 0$), population sizes can become unbounded due to the positive amplifying feedback loop between plants and pollinators that is no longer controlled by intra-specific competition. In such a case, if the plant population at carrying capacity is sufficient for the pollinator population to have a positive growth rate when low in density, then populations grow unbounded (mathematically converging toward infinity) irrespective of initial conditions. If not, the equilibrium where only plants are present is locally stable but populations grow unbounded if initially large enough.

### III. Ecological equilibria in the plant-pollinator-herbivore community

#### 1. Dynamical system in matrix form

The dynamical system capturing population dynamics (i.e. equations (a, b, c)) can be written in matrix form as:

$$\begin{pmatrix} \frac{dP}{dt} \\ \frac{dM}{dt} \\ \frac{dH}{dt} \end{pmatrix} = \begin{pmatrix} r_p \\ r_m \\ r_h \end{pmatrix} + \begin{pmatrix} -c_p & a_{pm} & -a_{ph} \\ e_m a_{pm} & -c_m & 0 \\ e_h a_{ph} & 0 & -c_h \end{pmatrix} \begin{pmatrix} P \\ M \\ H \end{pmatrix}$$

The corresponding Jacobian matrix is written:

$$Jac = \begin{pmatrix} r_p - 2c_p P + a_{pm} M - a_{ph} H & a_{pm} P & -a_{ph} P \\ e_m a_{pm} M & r_m - 2c_m M + e_m a_{pm} P & 0 \\ e_h a_{ph} H & 0 & r_h - 2c_h H + e_h a_{ph} P \end{pmatrix}$$

#### 2. Expressions of equilibria and feasibility

| Equilibrium | Expression | Feasibility |
|---|---|---|
| Absence | $P^* = 0\,;\ M^* = 0\,;\ H^* = 0$ | Yes |
| Plants | $P^* = {}^{r_p}/_{c_p}\ ;\ M^* = 0\,;\ H^* = 0$ | Yes |
| Pollinators | $P^* = 0\,;\ M^* = {}^{r_m}/_{c_m}\ ;\ H^* = 0$ | No |
| Herbivores | $P^* = 0\,;\ M^* = 0\,;\ H^* = {}^{r_h}/_{c_h}$ | No |
| Plants and pollinators | $P^*_{PM} = \dfrac{c_m r_p + a_{pm} r_m}{c_p c_m - e_m a_{pm}^2}$<br>$M^*_{PM} = \dfrac{e_m a_{pm} r_p + c_p r_m}{c_p c_m - e_m a_{pm}^2}\ ;\ H^*_{PM} = 0$ | $\begin{cases} c_p c_m - e_m a_{pm}^2 \geq 0 \\ r_m + e_m a_{pm}\left[\dfrac{r_p}{c_p}\right] \geq 0 \end{cases}$<br>Or<br>$\begin{cases} c_p c_m - e_m a_{pm}^2 \leq 0 \\ r_m + e_m a_{pm}\left[\dfrac{r_p}{c_p}\right] \leq 0 \end{cases}$ |
| Plants and herbivores | $P^*_{PH} = \dfrac{c_h r_p - a_{ph} r_h}{c_p c_h + e_h a_{ph}^2}$<br>$M^*_{PH} = 0\,;\ H^*_{PH} = \dfrac{e_h a_{ph} r_p + c_p r_h}{c_p c_h + e_h a_{ph}^2}$ | $r_h + e_h a_{ph}\left[\dfrac{r_p}{c_p}\right] \geq 0$ |
| Pollinators and herbivores | $P^*_{MH} = 0\,;\ M^*_{MH} = {}^{r_m}/_{c_m}\ ;\ H^*_{MH} = {}^{r_h}/_{c_h}$ | No |
| **Coexistence** of plants, pollinators and herbivores | $P^* = \dfrac{c_h c_m r_p + c_h a_{pm} r_m - c_m a_{ph} r_h}{c_h c_m c_p - c_h e_m a_{pm}{}^2 + c_m e_h a_{ph}{}^2}$<br>$M^* = \dfrac{c_h e_m a_{pm} r_p + (c_p c_h + e_h a_{ph}{}^2) r_m - e_m a_{pm} a_{ph} r_h}{c_h c_m c_p - c_h e_m a_{pm}{}^2 + c_m e_h a_{ph}{}^2}$<br>$H^* = \dfrac{c_m e_h a_{ph} r_p + e_h a_{pm} a_{ph} r_m + (c_p c_m - e_m a_{pm}{}^2) r_h}{c_h c_m c_p - c_h e_m a_{pm}{}^2 + c_m e_h a_{ph}{}^2}$ | |

**Table S1: Expressions of equilibria and feasibility.** Reminder of our assumptions: $r_p > 0, r_m < 0, r_h < 0$

#### 3. Stability of equilibria

An equilibrium is stable if, and only if, the three eigenvalues of the Jacobian matrix, calculated at this equilibrium, have a negative real part. Such property is equivalent to the three Routh-Hurwitz inequalities, which are often easier to assess. Note that we only assess the stability of the five potentially feasible equilibria.

| Equilibrium | Method | Mathematical expressions | Stability |
|---|---|---|---|
| Absence | Eigenvalues $(\lambda_1, \lambda_2, \lambda_3)$ | $\lambda_1 = r_p$ ; $\lambda_2 = r_m$ ; $\lambda_3 = r_h$ | No |
| Plants | Eigenvalues $(\lambda_1, \lambda_2, \lambda_3)$ | $\lambda_1 = -r_p$<br>$\lambda_2 = \dfrac{e_m a_{pm} r_p + c_p r_m}{c_p}$<br>$\lambda_3 = \dfrac{e_h a_{ph} r_p + c_p r_h}{c_p}$ | $\begin{cases} r_m + e_m a_{pm} \left[\dfrac{r_p}{c_p}\right] \leq 0 \\ r_h + e_h a_{ph} \left[\dfrac{r_p}{c_p}\right] \leq 0 \end{cases}$ |
| Plants and pollinators | Routh-Hurwitz | $a_1 = \left(c_p - e_h a_{ph}\right) P^*_{PM} + c_m M^*_{PM} - r_h$<br>$a_2 = P^*_{PM} M^*_{PM} \left(c_p c_m - e_h c_m a_{ph} - e_m a^2_{pm}\right) - M^*_{PM} c_m r_h - c_p P^*_{PM} (e_h a_{ph} P^*_{PM} + r_h)$<br>$a_3 = P^*_{PM} M^*_{PM} (e_h a_{ph} P^*_{PM} + r_h)(e_m a^2_{pm} - c_p c_m)$ | |
| Plants and herbivores | Routh-Hurwitz | $a_1 = \left(c_p - a_{pm} e_m\right) P^*_{PH} + c_h H^*_{PH} - r_m$<br>$a_2 = P^*_{PH} H^*_{PH} \left(c_h c_p - a_{pm} c_h e_m + a^2_{ph} e_h\right) - H^*_{PH} c_h r_m - c_p P^*_{PH} (a_{pm} e_m P^*_{PH} + r_m)$<br>$a_3 = -P^*_{PH} H^*_{PH} (e_m a_{pm} P^*_{PH} + r_m)(e_h a^2_{ph} + c_h c_p)$ | |
| **Coexistence** of plants, pollinators and herbivores | Routh-Hurwitz | $a_1 = c_p P^* + c_m M^* + c_h H^*$<br>$a_2 = P^* M^* \left(c_p c_m - e_m a^2_{pm}\right) + P^* H^* \left(c_p c_h + e_h a^2_{ph}\right) + M^* H^* c_m c_h$<br>$a_3 = P^* M^* H^* (c_h c_m c_p - c_h e_m {a_{pm}}^2 + c_m e_h {a_{ph}}^2)$ | |

**Table S2: Mathematical expressions of local stability.** A necessary and sufficient condition for stability is either: $(\lambda_1 < 0; \lambda_2 < 0; \lambda_3 < 0)$ or $(a_1 > 0;\ a_3 > 0;\ a_1 a_2 - a_3 > 0)$. Reminder of our assumptions: $r_p > 0, r_m < 0, r_h < 0$

### IV. First analytical results

#### 1. Variation of biomass densities with interaction strengths

Plant density at stable coexistence increases with pollination and decreases with herbivory.

We have:

$$P^* = \frac{c_h c_m r_p + c_h a_{pm} r_m - c_m a_{ph} r_h}{c_h c_m c_p - c_h e_m a_{pm}{}^2 + c_m e_h a_{ph}{}^2}$$

$$\Leftrightarrow (c_h c_m c_p - c_h e_m a_{pm}{}^2 + c_m e_h a_{ph}{}^2)P^* = c_h c_m r_p + c_h a_{pm} r_m - c_m a_{ph} r_h$$

$$\Rightarrow \begin{cases} -2c_h e_m a_{pm} P^* + (c_h c_m c_p - c_h e_m a_{pm}{}^2 + c_m e_h a_{ph}{}^2)\dfrac{\partial P^*}{\partial a_{pm}} = c_h r_m \\ 2c_m e_h a_{ph} P^* + (c_h c_m c_p - c_h e_m a_{pm}{}^2 + c_m e_h a_{ph}{}^2)\dfrac{\partial P^*}{\partial a_{ph}} = -c_m r_h \end{cases}$$

$$\Rightarrow \begin{cases} \dfrac{\partial P^*}{\partial a_{pm}} = \dfrac{c_h(r_m + 2e_m a_{pm} P^*)}{(c_h c_m c_p - c_h e_m a_{pm}{}^2 + c_m e_h a_{ph}{}^2)} \\ \dfrac{\partial P^*}{\partial a_{ph}} = \dfrac{-c_m(r_h + 2e_h a_{ph} P^*)}{(c_h c_m c_p - c_h e_m a_{pm}{}^2 + c_m e_h a_{ph}{}^2)} \end{cases}$$

$$\Rightarrow \begin{cases} \dfrac{\partial P^*}{\partial a_{pm}} = \dfrac{c_h(c_m M^* + e_m a_{pm} P^*)}{(c_h c_m c_p - c_h e_m a_{pm}{}^2 + c_m e_h a_{ph}{}^2)} \\ \dfrac{\partial P^*}{\partial a_{ph}} = \dfrac{-c_m(c_h H^* + e_h a_{ph} P^*)}{(c_h c_m c_p - c_h e_m a_{pm}{}^2 + c_m e_h a_{ph}{}^2)} \end{cases}$$

$$\Rightarrow \begin{cases} \dfrac{\partial P^*}{\partial a_{pm}} > 0 \\ \dfrac{\partial P^*}{\partial a_{ph}} < 0 \end{cases}$$

Given that $P^*, M^*, H^* > 0$ (feasibility) and $(c_h c_m c_p - c_h e_m a_{pm}{}^2 + c_m e_h a_{ph}{}^2) > 0$ (stability).

Moreover,

$$c_m M^* = r_m + e_m a_{pm} P^*$$

$$\Rightarrow \begin{cases} \dfrac{\partial M^*}{\partial a_{pm}} = \dfrac{e_m}{c_m}\left(P^* + a_{pm}\dfrac{\partial P^*}{\partial a_{pm}}\right) > 0 \\ \dfrac{\partial M^*}{\partial a_{ph}} = \dfrac{e_m}{c_m} a_{pm} \dfrac{\partial P^*}{\partial a_{ph}} > 0 \end{cases}$$

Pollinator density at stable coexistence increases with pollination and decreases with herbivory.

And

$$c_h H^* = r_h + e_h a_{ph} P^*$$

$$\Rightarrow \begin{cases} \dfrac{\partial H^*}{\partial a_{pm}} = \dfrac{e_h}{c_h} a_{ph} \dfrac{\partial P^*}{\partial a_{pm}} > 0 \\ \dfrac{\partial H^*}{\partial a_{ph}} = \dfrac{e_h}{c_h}\left(P^* + a_{ph}\dfrac{\partial P^*}{\partial a_{ph}}\right) = e_h \dfrac{P^*(c_m c_p - e_m a_{pm}{}^2) - c_m a_{ph} H^*}{(c_h c_m c_p - c_h e_m a_{pm}{}^2 + c_m e_h a_{ph}{}^2)} \end{cases}$$

Herbivore density increases with pollination. It can either increase or decrease with herbivory. It necessarily decreases if pollination is strong ($e_m a_{pm}{}^2 > c_m c_p$) or herbivore density ($H^*$) is high.

#### 2. Global stability of the coexistence equilibrium when unbounded growth is not possible

We prove analytically that when populations cannot become unbounded in the plant-pollinator subcommunity, the feasibility of the coexistence equilibrium implies its global stability.

*Proof of the global stability of feasible coexistence in the case ($c_p c_m - e_m a_{pm}^2 > 0$)

The nondimensionalized dynamical system (Appendix A) is rewritten in matrix form:

$$\begin{pmatrix} \dfrac{dP}{dt} \\ \dfrac{dM}{dt} \\ \dfrac{dH}{dt} \end{pmatrix} = \begin{pmatrix} 1 \\ \rho_m \\ \rho_h \end{pmatrix} + A \begin{pmatrix} P \\ M \\ H \end{pmatrix}$$

$$A = \begin{pmatrix} -1 & \alpha_{pm} & -\alpha_{ph} \\ \lambda_m \alpha_{pm} & -1 & 0 \\ \lambda_h \alpha_{ph} & 0 & -1 \end{pmatrix}$$

Let D be the positive diagonal matrix defined as:

$$D = \begin{pmatrix} 1 & 0 & 0 \\ 0 & \dfrac{1}{\lambda_m} & 0 \\ 0 & 0 & \dfrac{1}{\lambda_h} \end{pmatrix}$$

We have:

$$D(-A)+(-A^t)D=\begin{pmatrix}1 & -\alpha_{pm} & \alpha_{ph}\\ -\alpha_{pm} & \dfrac{1}{\lambda_m} & 0\\ -\alpha_{ph} & 0 & \dfrac{1}{\lambda_h}\end{pmatrix}+\begin{pmatrix}1 & -\alpha_{pm} & \alpha_{ph}\\ -\alpha_{pm} & \dfrac{1}{\lambda_m} & 0\\ -\alpha_{ph} & 0 & \dfrac{1}{\lambda_h}\end{pmatrix}=\begin{pmatrix}2 & -2\alpha_{pm} & 0\\ -2\alpha_{pm} & \dfrac{2}{\lambda_m} & 0\\ 0 & 0 & \dfrac{2}{\lambda_h}\end{pmatrix}$$

$D(-A)+(-A^t)D$ is positive definite since it satisfies the Silvester Criterion:

(1) $|2| = 2 > 0$

(2) $\begin{vmatrix}2 & -2\alpha_{pm}\\ -2\alpha_{pm} & \frac{2}{\lambda_m}\end{vmatrix} = \frac{4}{\lambda_m}\left(1-\lambda_m\alpha_{pm}^2\right) > 0$

(3) $\begin{vmatrix}2 & -2\alpha_{pm} & 0\\ -2\alpha_{pm} & \frac{2}{\lambda_m} & 0\\ 0 & 0 & \frac{2}{\lambda_h}\end{vmatrix} = \frac{2}{\lambda_h}\begin{vmatrix}2 & -2\alpha_{pm}\\ -2\alpha_{pm} & \frac{2}{\lambda_m}\end{vmatrix} = \frac{8}{\lambda_m\lambda_h}\left(1-\lambda_m\alpha_{pm}^2\right) > 0$

This implies that the matrix $A$ is VL-stable (Hofbauer *et al.* 2008) and in turn, that if the coexistence equilibrium is strictly feasible $(P^* > 0, M^* > 0, H^* > 0)$, it is globally stable (Logofet 2005; Hofbauer *et al.* 2008). All trajectories starting with strictly positive population densities converge toward this equilibrium.

Moreover, if the coexistence equilibrium is not strictly feasible, either the plant-pollinator or plant-herbivore equilibrium is feasible and globally stable. If not, the plant equilibrium is feasible and globally stable. The equilibrium that is feasible and globally stable attracts all solutions starting from strictly positive population densities (Hofbauer et al., 2008).

### Appendix C: Relationships governing stable coexistence

#### I. Characterization of stable coexistence

##### 1. Feasibility of the coexistence equilibrium

The feasibility corresponds to:

$$\begin{cases} P^* \geq 0 \\ M^* \geq 0 \\ H^* \geq 0 \end{cases}$$

However, the plant population supports the animal populations in our community ($r_m < 0, r_h < 0$) so that as long as at least one animal population is viable, the plant population is necessarily viable:

$$M^* \geq 0 \; or \; H^* \geq 0 \Rightarrow P^* \geq 0$$

<u>Proof:</u>
The coexistence equilibrium satisfies:

$$\begin{cases} P^* = \dfrac{-r_m + c_m M^*}{e_m a_{pm}} \geq \dfrac{-r_m}{e_m a_{pm}} > 0 \\ P^* = \dfrac{-r_h + c_h H^*}{e_h a_{ph}} \geq \dfrac{-r_h}{e_h a_{ph}} > 0 \end{cases}$$

Hence:

$$Feasability \Leftrightarrow \begin{cases} M^* \geq 0 \\ H^* \geq 0 \end{cases}$$

Moreover

$$M^* = \frac{c_h e_m a_{pm} r_p + (c_p c_h + e_h a_{ph}{}^2) r_m - e_m a_{pm} a_{ph} r_h}{c_h c_m c_p - c_h e_m a_{pm}{}^2 + c_m e_h a_{ph}{}^2}$$

$$= \frac{(c_p c_h + e_h a_{ph}{}^2)(r_m + e_m a_{pm} \overbrace{\dfrac{c_h r_p - a_{ph} r_h}{c_p c_h + e_h a_{ph}^2}}^{P^*_{PH}})}{c_h c_m c_p - c_h e_m a_{pm}{}^2 + c_m e_h a_{ph}{}^2}$$

$$H^* = \frac{c_m e_h a_{ph} r_p + e_h a_{pm} a_{ph} r_m + (c_p c_m - e_m a_{pm}{}^2) r_h}{c_h c_m c_p - c_h e_m a_{pm}{}^2 + c_m e_h a_{ph}{}^2}$$

$$= \frac{(c_p c_m - e_m a_{pm}{}^2)(r_h + e_h a_{ph} \overbrace{\dfrac{c_m r_p + a_{pm} r_m}{c_p c_m - e_m a_{pm}^2}}^{P^*_{PM}})}{c_h c_m c_p - c_h e_m a_{pm}{}^2 + c_m e_h a_{ph}{}^2}$$

Given than the denominator $c_h c_m c_p - c_h e_m {a_{pm}}^2 + c_m e_h {a_{ph}}^2$ can either be positive or negative, the coexistence equilibrium is feasible if and only if:

$$\begin{cases} c_h c_m c_p - c_h e_m {a_{pm}}^2 + c_m e_h {a_{ph}}^2 > 0 \\ (c_p c_h + e_h {a_{ph}}^2)(r_m + e_m a_{pm} P_{PH}^*) \geq 0 \\ (c_p c_m - e_m {a_{pm}}^2)(r_h + e_h a_{ph} P_{PM}^*) \geq 0 \end{cases}$$

Or

$$\begin{cases} c_h c_m c_p - c_h e_m {a_{pm}}^2 + c_m e_h {a_{ph}}^2 < 0 \\ (c_p c_h + e_h {a_{ph}}^2)(r_m + e_m a_{pm} P_{PH}^*) \leq 0 \\ (c_p c_m - e_m {a_{pm}}^2)(r_h + e_h a_{ph} P_{PM}^*) \leq 0 \end{cases}$$

#### 2. Stability of the coexistence equilibrium

Given it is feasible, the coexistence equilibrium is stable if and only if the three Routh-Hurwitz conditions are satisfied:

$$c_p P^* + c_m M^* + c_h H^* > 0$$

$$P^* M^* H^* \left(c_h c_m c_p - c_h e_m {a_{pm}}^2 + c_m e_h {a_{ph}}^2\right) > 0$$

$$\left(c_p P^* + c_m M^* + c_h H^*\right)\left(P^* M^* \left(c_p c_m - e_m a_{pm}^2\right) + P^* H^* \left(c_p c_h + e_h a_{ph}^2\right) + M^* H^* c_m c_h\right) - P^* M^* H^* \left(c_h c_m c_p - c_h e_m {a_{pm}}^2 + c_m e_h {a_{ph}}^2\right) > 0$$

The first condition is always true (feasibility). Stability is therefore characterized by the two following relationships, which correspond respectively to relationships (3) and (4) in the main text (table 2).

$$\left(c_h c_m c_p - c_h e_m {a_{pm}}^2 + c_m e_h {a_{ph}}^2\right) > 0$$

$$\left(c_p P^* + c_m M^* + c_h H^*\right)\left(P^* M^* \left(c_p c_m - e_m a_{pm}^2\right) + P^* H^* \left(c_p c_h + e_h a_{ph}^2\right) + M^* H^* c_m c_h\right) - P^* M^* H^* \left(c_h c_m c_p - c_h e_m {a_{pm}}^2 + c_m e_h {a_{ph}}^2\right) > 0$$

### II. Relationships governing stable coexistence

By combining the inequations characterizing feasibility and stability, we obtain a system of four inequations that are necessary and sufficient for the stable coexistence of plants, pollinators, and herbivores within our framework:

$$\begin{cases} c_h c_m c_p - c_h e_m {a_{pm}}^2 + c_m e_h {a_{ph}}^2 > 0 \\ (c_p c_h + e_h {a_{ph}}^2)(r_m + e_m a_{pm} P_{PH}^*) \geq 0 \\ (c_p c_m - e_m {a_{pm}}^2)(r_h + e_h a_{ph} P_{PM}^*) \geq 0 \\ \left(c_p P^* + c_m M^* + c_h H^*\right)\left(P^* M^* \left(c_p c_m - e_m a_{pm}^2\right) + P^* H^* \left(c_p c_h + e_h a_{ph}^2\right) \right. \\ \left. + M^* H^* c_m c_h\right) - P^* M^* H^* \left(c_h c_m c_p - c_h e_m {a_{pm}}^2 + c_m e_h {a_{ph}}^2\right) > 0 \end{cases}$$

$$\Leftrightarrow \begin{cases} c_h c_m c_p - c_h e_m {a_{pm}}^2 + c_m e_h {a_{ph}}^2 > 0 \\ r_m + e_m a_{pm} P_{PH}^* \geq 0 \\ (c_p c_m - e_m {a_{pm}}^2)(r_h + e_h a_{ph} P_{PM}^*) \geq 0 \\ \left(c_p P^* + c_m M^* + c_h H^*\right)\left(P^* M^* \left(c_p c_m - e_m a_{pm}^2\right) + P^* H^* \left(c_p c_h + e_h a_{ph}^2\right) \right. \\ \left. + M^* H^* c_m c_h\right) - P^* M^* H^* \left(c_h c_m c_p - c_h e_m {a_{pm}}^2 + c_m e_h {a_{ph}}^2\right) > 0 \end{cases}$$

These relationships correspond respectively to the relationships (1), (2), (3), and (4) in the main text (table 2). In what follows, we present for each relationship how its biological interpretation, as well as the influence of each type of interaction, are derived (3rd and 4th columns of table 2).

#### 1. Relationship (1)

| | |
|---|---|
| $r_m + e_m a_{pm} P_{PH}^* \geq 0$ <br> $P_{PH}^* \overset{\text{def}}{=} \dfrac{c_h r_p - a_{ph} r_h}{c_p c_h + e_h a_{ph}^2}$ | (1) |

Assuming that the plant-herbivore equilibrium is feasible, we have:

$$\left.\frac{1}{M}\frac{dM}{dt}\right)_{M \to 0^+} = r_m + e_m a_{pm} P_{PH}^*$$

Relationship (1) consequently means that the per capita growth rate of pollinators population, when low in density, is positive within the plant-herbivore community at ecological equilibrium. Thus, pollinators are able to invade the plant-herbivore community. This biological interpretation is valid as long as the plant-herbivore equilibrium is feasible.

Moreover, given that $P_{PH}^*$ is always positive, relationship (1) is equivalent to relationship (1’):

| | |
|---|---|
| $\dfrac{-r_m}{e_m P_{PH}^*} \leq a_{pm}$ | (1′) |

*We can study the variations of $P_{PH}^*$ with $a_{ph}$ (using the nondimensionalized system for simplicity):

$$(1+\lambda_h\alpha_{ph}^2)^2\frac{\partial P_{PH}^*}{\partial\alpha_{ph}} = \rho_h\lambda_h\alpha_{ph}^2 - 2\lambda_h\alpha_{ph} - \rho_h$$

$$\Leftrightarrow \left(1+\lambda_h\alpha_{ph}^2\right)\frac{\partial P_{PH}^*}{\partial\alpha_{ph}} = -\lambda_h\alpha_{ph}P_{PH}^* - H_{PH}^*$$

The latter first indicates that as long as the plant-herbivore equilibrium is feasible (i.e. $a_{ph} \geq \frac{-c_p r_h}{e_{hrp}}$), $P_{PH}^*$ decreases with herbivory. Moreover, the polynomial on the right side of the first-line equality (i.e. $\rho_h\lambda_h\alpha_{ph}^2 - 2\lambda_h\alpha_{ph} - \rho_h$) has two roots ($root1_\pm = \frac{1\pm\sqrt{1+\rho_h^2/\lambda_h}}{\rho_h}$), only one ($root1_-$) being positive. When at this root, the polynomial switches from positive to negative indicating that $P_{PH}^*$ is maximal. Relationship (1) can therefore never be satisfied if pollination is below a critical value:

$$\alpha_{pm} < \alpha_{pm}^{critical} \stackrel{\text{def}}{=} \frac{2\lambda_h\rho_m}{\lambda_m\rho_h^2}\left(1-\sqrt{1+{\rho_h^2}/{\lambda_h}}\right)$$

The behavior of relationship (1) is now fully determined. **Table S3** recapitulates the analysis in the dimensionalized language. It notably shows that relationship (1) behaves according to biological intuition (i.e. $P_{PH}^*$ decreases with herbivory) over the domain of feasibility of the plant-herbivore equilibrium (**green, table S3**).

| Variation of $P_{PH}^*$ with herbivory ($a_{ph}$) | $a_{ph}$ | $0$ | $\frac{c_h r_p}{r_h}\left[1-\sqrt{1+\frac{c_p r_h^2}{e_h c_h r_p^2}}\right]$ | $\frac{-c_p r_h}{e_h r_p}$ | $+\infty$ |
|---|---|---|---|---|---|
| | $P_{PH}^*$ | $\frac{r_p}{c_p}$ ↗ | $\max(P_{PH}^*)$ | ↘ | $0$ |

**Table S3: Analytical study ore relationship (1).** The biological interpretation of relationship (1) is valid over the domain of feasibility of the plant-herbivore equilibrium (green).

#### 2. Relationship (2)

| | |
|---|---|
| $(c_p c_m - e_m a_{pm}{}^2)(r_h + e_h a_{ph} P_{PM}^*) \geq 0$<br>$P_{PM}^* \stackrel{\text{def}}{=} \frac{c_m r_p + a_{ph} r_m}{c_p c_m - e_m a_{pm}^2}$ | (2) |

Let us consider three quantities of interest here $a_{pm}^{M} \stackrel{\text{def}}{=} \frac{-r_m c_p}{e_m r_p}$ ; $a_{pm}^{P} \stackrel{\text{def}}{=} \frac{-c_m r_p}{r_m}$ ; $a_{pm}^{\infty} \stackrel{\text{def}}{=} \sqrt{\frac{c_p c_m}{e_m}}$ :

- $a_{pm}^{M}$ is the pollination level above which pollinators are able to invade the plant community when low in density; it is also the value of pollination at which the numerator of $M_{PM}^{*}$ switches sign, from negative to positive

- $a_{pm}^{P}$ is the value of pollination at which the numerator of $P_{PM}^{*}$ switches sign, from positive to negative

- $a_{pm}^{\infty}$ is the pollination level above which unbounded population growth becomes possible; it is also the value of pollination at which the denominator of $P_{PM}^{*}$ and $M_{PM}^{*}$ switches sign, from positive to negative

There are only two ways of ordering these three pollination levels, which are illustrated in **Fig. S1 and S2.** We invite the reader to refer to these figures in order to facilitate the comprehension of the subsequent mathematical analysis. Also note that an important distinction between the two cases is the occurrence of unbounded growth depending on initial densities in case (b), a feature enabling the occurrence of alternative stable states at the three-species community scale.

- <u>Case (1):</u> $a_{pm}^{M} < a_{pm}^{\infty} < a_{pm}^{P}$ (this case corresponds to the parameter set of Fig. 2&3)

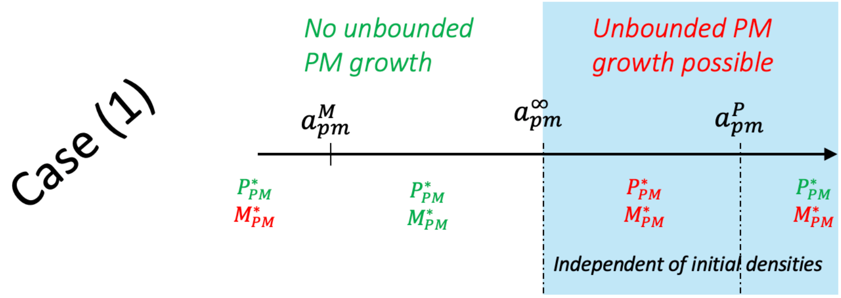


**Fig. S1: Schematic representation of the parameter instance of case (1).** Densities in red are negative; densities in green are positive.

- <u>Case (2):</u> $a_{pm}^{P} < a_{pm}^{\infty} < a_{pm}^{M}$ (this case corresponds to the parameter set of Fig. S3)

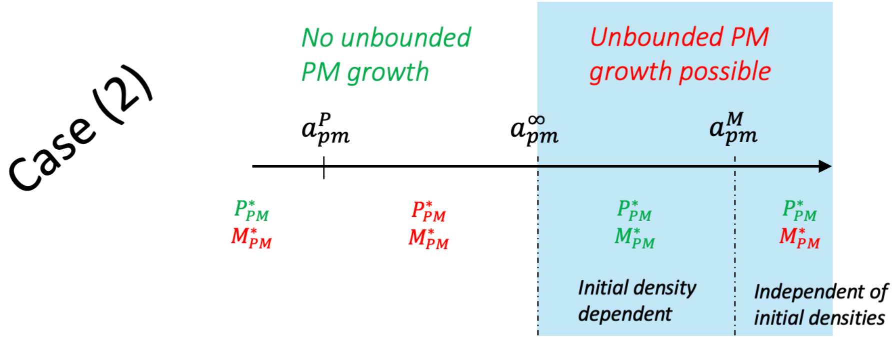


**Fig. S2: Schematic representation of the parameter instance of case (2).** Densities in red are negative; densities in green are positive.

$P^*_{PM}$ is negative for pollination between $min\,(a^P_{pm}, a^\infty_{pm})$ and $max\,(a^P_{pm}, a^\infty_{pm})$. Therefore, relationship (2) is always satisfied in case (1) for pollination in the interval $[a^\infty_{pm}, a^P_{pm}]$; while it is never satisfied in case (2) for pollination in the interval $[a^P_{pm}, a^\infty_{pm}]$. When pollination is not between $min\,(a^P_{pm}, a^\infty_{pm})$ and $max\,(a^P_{pm}, a^\infty_{pm})$, because $P^*_{PM}$ is positive, relationship (2) can take two forms according to the sign of $c_p c_m - e_m {a_{pm}}^2$. As in the main document, these two forms are labeled $(2'a)$ and $(2'b)$ (see the last row of **Table S4**).

| | |
|---|---|
| $\begin{cases} a_{pm} \le a^\infty_{pm} \\ \dfrac{-r_h}{e_h P^*_{PM}} \le a_{ph} \end{cases}$ | $(2'a)$ |
| $\begin{cases} a_{pm} \ge a^\infty_{pm} \\ a_{ph} \le \dfrac{-r_h}{e_h P^*_{PM}} \end{cases}$ | $(2'b)$ |

Given that $\frac{-r_h}{e_h P^*_{PM}} \le a_{ph} \Leftrightarrow (r_h + e_h a_{ph} P^*_{PM}) \stackrel{\text{def}}{=} \left.\frac{1}{H}\frac{dH}{dt}\right)_{H\to 0^+} \ge 0$, relationship (2'a) corresponds biologically to the herbivores being able to invade the plant-pollinator community at ecological equilibrium, when low in density and unbounded PM growth is not possible. Note that this interpretation (**Table 2, (2a)**) requires the plant-pollinator equilibrium to be feasible. Relationship (2'b) sets an upper limit to the intensity of herbivory when unbounded PM growth is possible. We interpret such a limit as follows: herbivores cannot consume the plant resource too strongly, otherwise, the plant resource density becomes too low for pollinators to persist (**Table 2, (2b)**).

Let us now study the variations of $P^*_{PM}$ with pollination ($a_{pm}$). Just before that, we show that the coexistence equilibrium is not possible below the $min\,(a^P_{pm}, a^M_{pm})$ so that we can exclude the corresponding interval from our study of variations.

Proof:

When $a_{pm} \le min\,(a^P_{pm}, a^M_{pm})$, we necessarily have that $a_{pm} \le a^\infty_{pm}$. Consequently, the plant equilibrium is globally stable within the PM subsystem (because unbounded growth is not possible). Thus, all trajectories of the PM dynamical subsystem (population dynamics, equations (a)(b) with $a_{ph} = 0$ ) starting with strictly positive initial conditions converge toward the plant equilibrium.

Now assume, by the absurd, that the plant-pollinator-herbivore coexistence equilibrium is strictly feasible. Because unbounded growth is not possible, the coexistence equilibrium is globally stable: all trajectories of the dynamical system (population dynamics, equations (a)(b)(c)) starting with strictly positive initial conditions converge toward the coexistence equilibrium.

Consider such a trajectory $\Psi_3$, its projection on the $(P, M)$ space ($\mathbb{R}^{+2}$) ($\Psi_2$) and the trajectory of the PM subsystem ($\Psi$), that has same initial conditions as the projection. $\Psi_2$ is a sub-solution of the PM dynamical system, so that we have that $\Psi$ is always above $\Psi_2$ (Hulin 2020). By letting time goes toward infinity (which we can do because both $\Psi$ and $\Psi_2$ converges toward finite values), we deduce that $M^* \stackrel{\text{def}}{=} \lim_{t\to+\infty} {\Psi_2}^{(2)} \leq \lim_{t\to+\infty} \Psi^{(2)} = 0$, which is in contradiction with the coexistence equilibrium being strictly feasible. ■

*Study of the variation of $P^*_{PM}$ with pollination $a_{pm}$:

By nondimensionalizing the system, we study the variations of $P^*_{PM} \stackrel{\text{def}}{=} \frac{1+\rho_m\alpha_{pm}}{1-\lambda_m\alpha^2_{pm}}$ with pollination ($\alpha_{pm}$). We remind the reader that $P^*_{PM}(a_{pm})$ is, first of all, a mathematical function. It biologically corresponds to the plant population at plant-pollinator equilibrium only with such an equilibrium is feasible ($P^*_{PM} > 0$ & $M^*_{PM} > 0$) and stable.

$$(1-\lambda_m\alpha^2_{pm})^2\frac{\partial P^*_{PM}}{\partial\alpha_{pm}} = \rho_m\lambda_m\alpha^2_{pm} + 2\lambda_m\alpha_{pm} + \rho_m$$

$$\Leftrightarrow \left(1-\lambda_m\alpha^2_{pm}\right)\frac{\partial P^*_{PM}}{\partial\alpha_{pm}} = \lambda_m\alpha_{pm}P^*_{PM} + M^*_{PM}$$

The last equation indicates that when the PM equilibrium is feasible, the plant density at this equilibrium increase with pollination below $a^{\infty}_{pm}$ and decreases above $a^{\infty}_{pm}$. In the latter case, $P^*_{PM}$ has to be regarded as a mathematical function, with no biological significance. Indeed, above $a^{\infty}_{pm}$, the model indicates that the plant biomass density is infinitely large ("orgy", unbounded growth). In other words, biological intuition does not apply since $P^*_{PM}$ does not correspond to the model output for plant density (but is rather a technical intermediate).

To go further, we study the polynomial that appears on the right side of the first equation (i.e. $\rho_m\lambda_m\alpha^2_{pm} + 2\lambda_m\alpha_{pm} + \rho_m$).

- In case (2) (which implies $\rho^2_m > \lambda_m$), this polynomial has no real roots so that it is always negative, which implies that the mathematical function ($P^*_{PM}$) decreases with pollination.

- In case (1) ($\rho^2_m < \lambda_m$), this polynomial has two positive roots ($root2_{\pm} = \frac{(1\pm\sqrt{(1-\rho^2_m/\lambda_m)})}{-\rho_m}$). Plant density ($P^*_{PM}$) increases with pollination when the pollination intensity is between these two roots (note that $root2_+ > root2_-$). We deduce that the interval of feasibility of the plant-pollinator equilibrium in this case (2) (i.e. $\left[a^M_{pm}, a^{\infty}_{pm}\right]$) is included in the interval $[c_m root2_-, c_m\, root2_+]$ because $P^*_{PM}$ should increase over it. Moreover, it is straightforward that $root2_+ > a^P_{pm}$ ($\stackrel{\text{def}}{=} c_m(-1/\rho_m)$).

The behavior of relationship (2) is now fully determined. **Table S4** recapitulates the analysis in the dimensionalized language.

#### Analytical study of relationship (2)

| Case | | | | | | | |
|---|---|---|---|---|---|---|---|
| Case (1) | Variation of $P_{PM}^*$ with pollination ($a_{pm}$) | $a_{pm}$ | $a_{pm}^M = \frac{-r_m c_p}{e_m r_p}$ | $a_{pm}^\infty = \sqrt{\frac{c_p c_m}{e_m}}$ | $a_{pm}^P = \frac{-c_m r_p}{r_m}$ | $root2_+ = \frac{-c_m r_p}{r_m}\left[1+\sqrt{1-\frac{c_p r_m^2}{e_m c_m r_p^2}}\right]$ | $+\infty$ |
| | | $P_{PM}^*$ | $\frac{r_p}{c_p}$ ↗ $+\infty$ | ‖ $-\infty$ ↗ | 0 ↗ | ↘ | 0 |
| | Form of relationship (2) | | (2'a) | Necessarily Satisfied | (2'b) | | |

| Case | | | | | | |
|---|---|---|---|---|---|---|
| Case (2) | Variation of $P_{PM}^*$ with pollination ($a_{pm}$) | $a_{pm}$ | $a_{pm}^P = \frac{-c_m r_p}{r_m}$ | $a_{pm}^\infty = \sqrt{\frac{c_p c_m}{e_m}}$ | $a_{pm}^M = \frac{-r_m c_p}{e_m r_p}$ | $+\infty$ |
| | | $P_{PM}^*$ | 0 ↘ | $-\infty$ ‖ $+\infty$ ↘ | | 0 |
| | Form of relationship (2) | | Necessarily Non-Satisfied | (2'b) | | |

**Table S4: Analytical study ore relationship (2).** The domain of feasibility of the plant-pollinator equilibrium is indicated in green.

In the main document, a situation corresponding to case (1) is presented in **Fig. 3**. Here (**Fig. S2**), we present a situation corresponding to case (2).

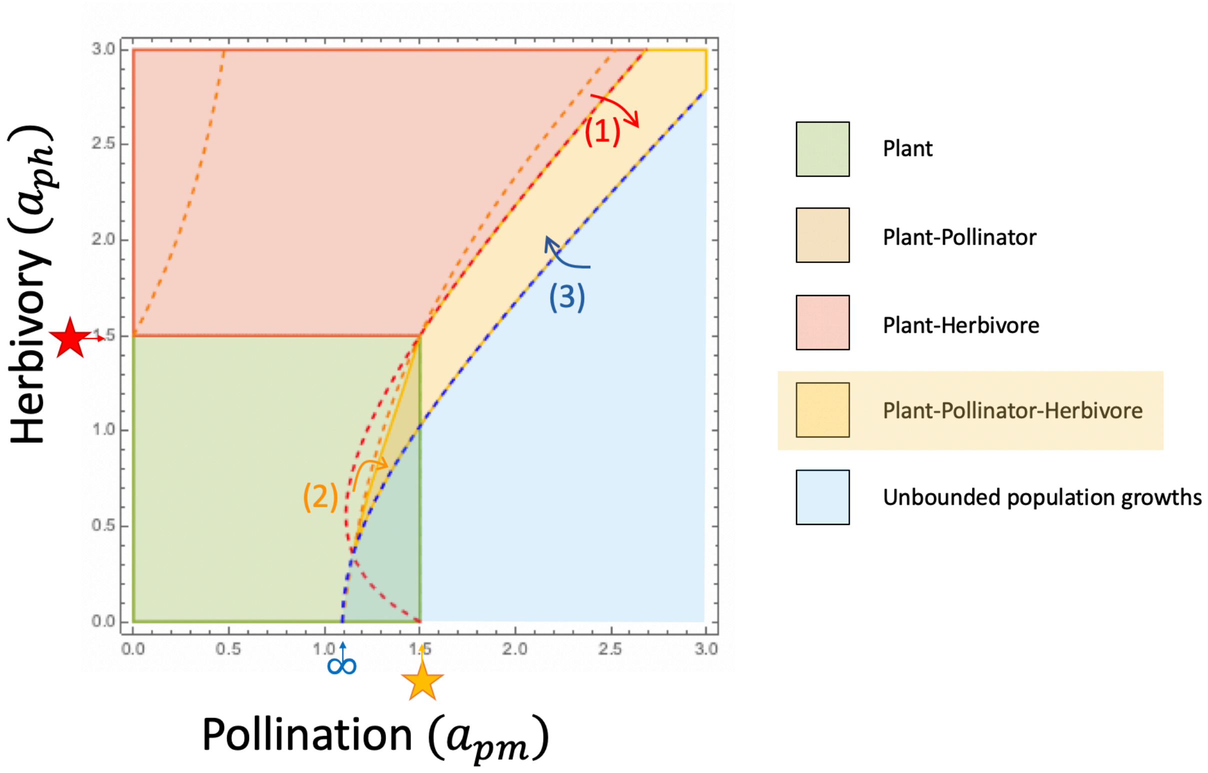


**Fig. S3: Community composition according to both pollination and herbivory.** In blue, no equilibrium is stable so that populations grow unboundedly. Arrows (1), (2), and (3) indicate the transitions enabling to satisfy relationships (1), (2), and (3) (**Table 2**), indicated by the dashed red, orange, and blue curve, respectively. These three relationships are not sufficient to achieve stable coexistence given the parameter set, indicating that relationship (4) is constraining here. More precisely, relationship (4) is the one that constrains the upper boundary of the stable coexistence area (yellow) between $a_{pm}^{\infty}$ (infinity symbol) and $a_{pm}^{M}$ (orange star). Moreover, for such pollination levels, relationship (2) takes the form (2'b). A red star indicates the level of herbivory enabling herbivores to invade the plant community. Note that stable coexistence (yellow area) requires the two interactions to be of similar magnitude.

**Fig. S3** permits to illustrate two important points:

- In case (2), the occurrence of alternative stable states is possible. Indeed, both the plant equilibrium and the coexistence equilibrium can be feasible and stable, when pollination is between $a_{pm}^{\infty}$ (symbol ∞, **Fig. S2**) and $a_{pm}^{M}$ (orange star, **Fig. S2**). The reason is that in the PM subcommunity, for such pollination levels, the plant equilibrium is locally stable but, if populations are initially large enough, they grow unboundedly. Given that the presence of herbivores can control such an unstable behavior when relationship (3) is satisfied, this leads to alternative stable states. Herbivory has nevertheless to be low enough so that the exploitation of the plant resource by herbivores does not bring the pollinator density in the catchment area of the plant equilibrium within the PM subsystem. The latter constraint corresponds to relationship (2), which sets an upper limit to the intensity of herbivory (relationship (2'b)).

- In **Fig. S3**, the upper limit of the stable coexistence area (yellow) when the strength of pollination is between $a_{pm}^{\infty}$ (symbol ∞) and $a_{pm}^{M}$ (orange star) is set by relationship (4).

##### 3. Relationship (3)

The total feedback at a given level $k$ is a summation of the strengths of all the feedback loops of length $k$ and that of all the combinations of disjunct (non-overlapping) feedback loops of shorter length containing $k$ elements (Neutel & Thorne 2014). Relationship (3) corresponds to the total feedback at level 3 being negative. Here, two of these loops are stabilizing because they are negative: the self-limitation competition loop $(-c_h c_m c_p)$ and the trophic loop $(-c_m e_h a_{ph}{}^2)$ while the mutualistic loop is positive and hence destabilizing $\left(c_h e_m a_{pm}{}^2\right)$.

##### 3. Relationship (4)

The quantity $F_1 \stackrel{\text{def}}{=} -(c_p P^* + c_m M^* + c_h H^*)$ corresponds to the feedback at level 1. The quantity $F_2 \stackrel{\text{def}}{=} -(P^* M^* \left(c_p c_m - e_m a_{pm}^2\right) + P^* H^* \left(c_p c_h + e_h a_{ph}^2\right) + M^* H^* c_m c_h)$ corresponds to the feedback at level 2. The quantity $F_3 \stackrel{\text{def}}{=} -P^* M^* H^* \left(c_h c_m c_p - c_h e_m a_{pm}{}^2 + c_m e_h a_{ph}{}^2\right)$ corresponds to the feedback at level 3. Relationship (4) can be written: $F_1 F_2 + F_3 > 0$. Given that $F_1 < 0$ and assuming relationship (3) (i.e. $F_3 < 0$), relationship (4) implies that $F_2$ is negative. Relationships (3) and (4) together imply that the feedback at each level is negative, which is a necessary condition for stability (Levins 1974). Therefore, relationship (4) can be interpreted as the fact that the negative feedback at level 3 (long time lag) is weaker than the product of negative feedback at lower levels (1 & 2, smaller time lag) (Levins 1974).

### Appendix D: The effect of the other parameters on stable coexistence

#### I. Animal intrinsic growth rates

##### 1. Effect on stable coexistence

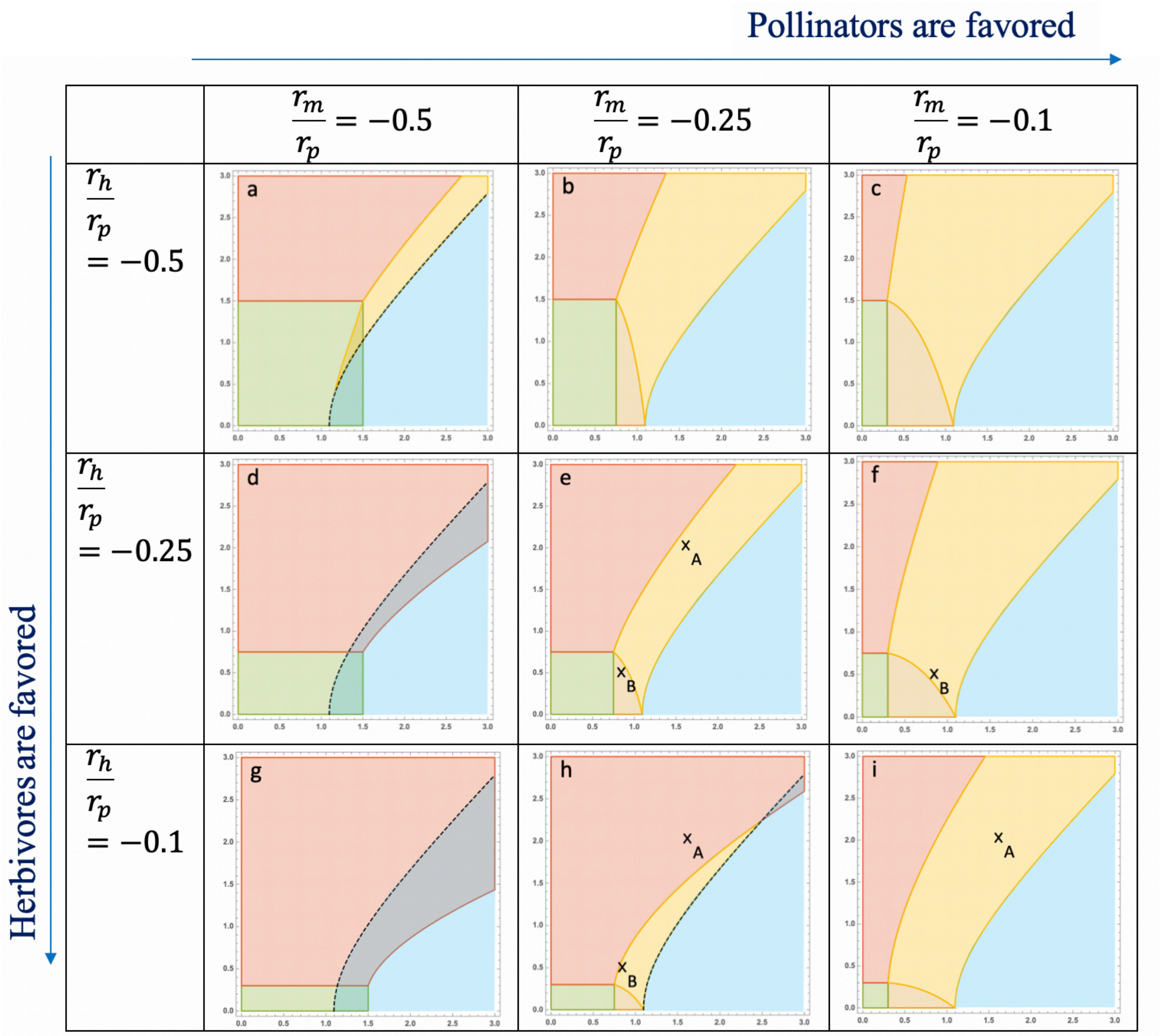


**Fig. S4: Effect of animal intrinsic growth rates on stable coexistence.** X-axis: Pollination ($a_{pm}$); Y-axis: Herbivory ($a_{ph}$) Colour legend: Green: Plant, Brown: Plant-Pollinator, Red: Plant-Herbivore, Yellow: stable coexistence, Light blue: unbounded population densities. In a-d-g-h, alternative (not both stable) states are observed (green and blue or red and blue overlap): if plant and pollinator densities are initially above a (herbivore-density-dependent) threshold, populations grow unboundedly; otherwise, pollinators are excluded. Note that **Fig. S4.a** corresponds to Fig. S3, characterized by the occurrence of alternative stable states (see Appendix C.II.2 for details). Parameters: $r_p = 10, e_m = e_h = 0.2,\ c_p = 0.6,\ c_m = c_h = 0.4$

Fig. 4A of the main document corresponds to Fig. S4 e-f-h-i. In addition to the analysis presented in the main text, we would like to highlight two points.

Firstly, stable coexistence is not possible if the pollinator intrinsic growth rate is too low in comparison with the herbivore one (Fig. S4 d&g). It is also much endangered if the intrinsic growth rates are too low, in spite of being similar (Fig. S4 a vs. i).

Secondly, for some parameter instances, unbounded population growth happens if the initial plant and pollinator densities are above a given threshold. This threshold gets higher as the initial herbivore density increases. In other words, depending on the initial control of the orgy of mutual benefaction by competition and herbivory, the community will display (or not) unbounded population dynamics. If the control is initially strong enough, pollinators are excluded. If herbivores are able to survive in the absence of pollinators, the final community consists of plants and herbivores at ecological equilibrium. If it is not the case, herbivores go extinct and only plants remain in the community.

#### 2. Analytical grounding

Animal intrinsic growth rates affect the feasibility of coexistence as indicated by relationships (1) and (2). We determine here how these relationships vary with the growth rates $r_m$ and $r_h$.

| | |
|---|---|
| $r_m + e_m a_{pm} P^*_{PH} \geq 0$ | (1) |

$$\frac{\partial(r_m + e_m a_{pm} P^*_{PH})}{\partial r_m} = 1 > 0$$

$$\frac{\partial(r_m + e_m a_{pm} P^*_{PH})}{\partial r_h} = e_m a_{pm} \frac{\partial P^*_{PH}}{\partial r_h} = \frac{-e_m a_{pm} a_{ph}}{c_p c_h + e_h a_{ph}^2} \leq 0$$

An increase in pollinator (resp. herbivore) growth rate facilitates (resp. impedes) the satisfaction of relationship (1).

| | |
|---|---|
| $(c_p c_m - e_m {a_{pm}}^2)(r_h + e_h a_{ph} P^*_{PM}) \geq 0$ | (2) |

$$\frac{\partial\left[(c_p c_m - e_m {a_{pm}}^2)(r_h + e_h a_{ph} P^*_{PM})\right]}{d r_m} = (c_p c_m - e_m {a_{pm}}^2) e_h a_{ph} \frac{\partial P^*_{PM}}{d r_m} = e_h a_{ph} a_{pm} \geq 0$$

$$\frac{\partial\left[(c_p c_m - e_m {a_{pm}}^2)(r_h + e_h a_{ph} P^*_{PM})\right]}{\partial r_h} = (c_p c_m - e_m {a_{pm}}^2)$$

An increase in pollinator growth rate makes it easier to satisfy relationship (2). An increase in herbivore growth rate makes it easier to satisfy relationship (2) when unbounded PM growth is not possible ($c_p c_m - e_m {a_{pm}}^2 > 0$), while it makes it harder when unbounded PM growth is possible.

### II. Animal intraspecific competition rates

#### 1. Effect on stable coexistence

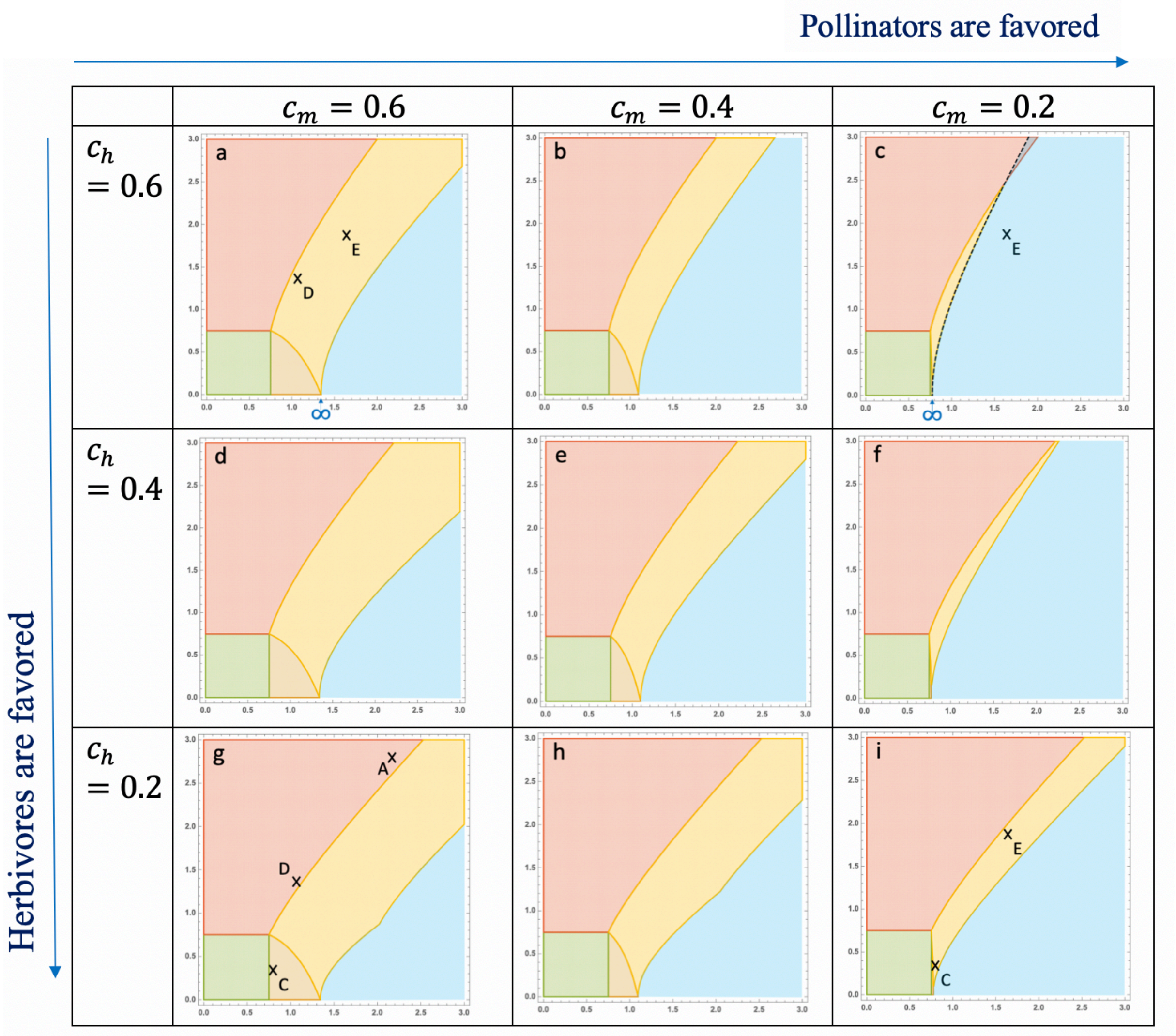


**Fig. S5: Effect of animal intraspecific competition on stable coexistence.** X-axis: Pollination ($a_{pm}$); Y-axis: Herbivory ($a_{ph}$) Colour legend: Green: Plant, Brown: Plant-Pollinator, Red: Plant-Herbivore, Yellow: stable coexistence, Light blue: unbounded population densities. In c, alternative states are observed (red and blue overlap): if plant and pollinator densities are initially above a (herbivore-density-dependent) threshold, populations grow unboundedly; otherwise, pollinators are excluded. In g & h, as pollination increases, the yellow-white border is first set by relationship (3) then (4), which explains the unusual shape. Parameters: $r_p = 10, r_m = r_h = -2.5, e_m = e_h = 0.2,\ c_p = 0.6$.

Fig. 4B of the main document corresponds to Fig. S5 a-c-g-i. In addition to the analysis presented in the main text, we would like to highlight one point. Feasibility mainly relies on the ability of a focal animal, at low population size and hence experimenting neglectable intraspecific competition, to invade the other animal-plant community (**Table 2**). As a result, the herbivore competition rate affects the pollinator ability to invade rather than its own (relationship (1)). Likewise, the pollinator competition rate affects the herbivore ability to invade rather than its own (relationship (2)).

#### 2. Analytical grounding

Animal intraspecific competitions affect both the feasibility and stability of coexistence (relationships (1) (2) (3) (4)). We determine here how relationships (1) (2) (3) vary with competition rates $c_m$ and $c_h$.

$$\frac{\partial(r_m + e_m a_{pm} P_{PH}^*)}{\partial c_m} = 0$$

$$\frac{\partial(r_m + e_m a_{pm} P_{PH}^*)}{\partial c_h} = e_m a_{pm} \frac{\partial P_{PH}^*}{\partial c_h} = e_m a_{pm} \frac{r_p\left(c_p c_h + e_h a_{ph}^2\right) - c_p(c_h r_p - a_{ph} r_h)}{(c_p c_h + e_h a_{ph}^2)^2}$$

$$= e_m a_{pm} \frac{a_{ph}(e_h a_{ph} r_p + c_p r_h)}{(c_p c_h + e_h a_{ph}^2)^2} = \frac{e_m a_{pm} a_{ph} H_{PH}^*}{c_p c_h + e_h a_{ph}^2}$$

Relationship (1) is not affected by competition among pollinators. When the plant-herbivore equilibrium is feasible, biological intuition applies: an increase in competition among herbivores makes it easier for pollinators to invade the plant-herbivore community due to an increase in plant density. When it is not, relationship (1) is impeded by an increase in herbivore competition rate.

$$\frac{\partial\left[(c_p c_m - e_m {a_{pm}}^2)(r_h + e_h a_{ph} P_{PM}^*)\right]}{\partial c_m} = (c_p c_m - e_m {a_{pm}}^2) e_h a_{ph} \frac{\partial P_{PM}^*}{\partial c_h} = -e_h a_{pm} a_{ph} M_{PM}^*$$

$$\frac{\partial\left[(c_p c_m - e_m {a_{pm}}^2)(r_h + e_h a_{ph} P_{PM}^*)\right]}{\partial c_h} = 0$$

Relationship (2) is not affected by competition among herbivores. When the plant-pollinator equilibrium is feasible, biological intuition applies: an increase in competition among pollinators makes it harder for herbivores to invade the plant-pollinator community due to a decrease in plant density. When it is not, relationship (2) is favored by an increase in pollinator competition rate.

| | |
|---|---|
| $c_h c_m c_p + c_m e_h {a_{ph}}^2 - c_h e_m {a_{pm}}^2 > 0$ | (3) |

$$\frac{\partial(c_h c_m c_p + c_m e_h {a_{ph}}^2 - c_h e_m {a_{pm}}^2)}{\partial c_m} = c_p c_h + e_h a_{ph}^2 > 0$$

$$\frac{\partial(c_h c_m c_p + c_m e_h {a_{ph}}^2 - c_h e_m {a_{pm}}^2)}{\partial c_h} = c_p c_m - e_m a_{pm}^2 < 0$$

The last inequality ensuing from the fact that it is meaningful to consider stability only when potential instability is possible (i.e. when $c_p c_m - e_m a_{pm}^2 < 0$), we conclude that an increase in pollinator (resp. herbivore) intraspecific competition favors (resp. disfavors) stability.

Finally, we can compare the effects on feasibility and on stability by approximating their ratio when interactions are very strong (i.e. at infinity). We find that stability effects overcome feasibility effects.

$$\frac{Feasibility\ effects}{Stability\ effects} = \frac{e_m a_{pm} a_{ph}(e_h a_{ph} r_p + c_p r_h)}{(c_p c_h + e_h a_{ph}^2)^2 (c_p c_m - e_m a_{pm}^2)} \sim \frac{r_p e_m e_h a_{pm} a_{ph}^2}{-e_m e_h^2 a_{pm}^2 a_{ph}^4} \sim -\frac{r_p}{e_h a_{pm} a_{ph}^2} \underbrace{\rightarrow}_{\substack{a_{pm}\rightarrow+\infty \\ a_{ph}\rightarrow+\infty}} 0$$

### III. Animal conversion efficiencies

#### 1. Effect on stable coexistence

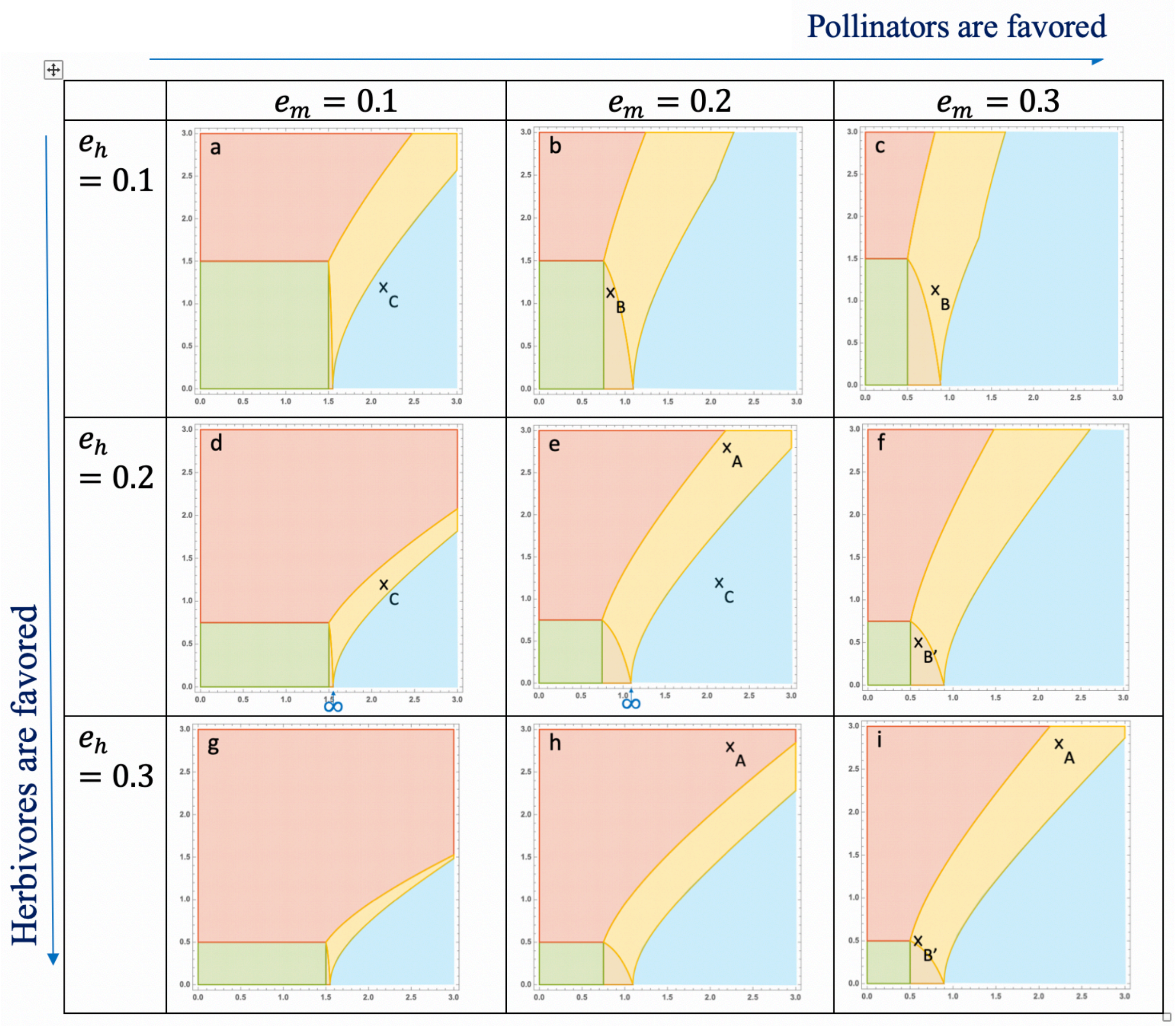


**Fig. S6: Effect of conversion efficiencies on stable coexistence.** X-axis: Pollination ($a_{pm}$); Y-axis: Herbivory ($a_{ph}$) Colour legend: Green: Plant, Brown: Plant-Pollinator, Red: Plant-Herbivore, Yellow: stable coexistence, Light blue: unbounded population densities. In b & c, as pollination increases, the yellow-white border is first set by relationship (3) then (4), which explains the unusual shape. Parameters: $r_p = 10, r_m = r_h = -2.5,\ c_p = 0.6,$ $c_m = c_h = 0.4$.

Overall, the range of pollination and herbivory intensities allowing stable coexistence gets wider in the upper right of **Fig. S6**, when pollinators exploit the plant resource more efficiently than herbivores. This pattern suggests that the effect of conversion efficiencies on feasibility (relationships (1), (2)) is stronger than their effect on stability (relationship (3)). As pollinators get more efficient at exploiting the plant resource, it becomes easier for them to invade the plant-herbivore community (**Fig. S6h vs S6i, point A**). It also makes it easier for herbivores to invade the plant-pollinator community as a result of an increased plant density (**Fig. S6b vs S6c, point B**). The positive feedback loop driven by pollination, however, gets stronger, favoring the orgy of mutual benefaction (**Fig. S6d vs S6e, point C and infinity symbol**). The effect of herbivores getting more efficient at exploiting their plant resource is contrasted on feasibility. On the one hand, it is easier for them to invade the plant-pollinator community (**Fig. S6f vs S6i, point B'**). On the other hand, an increased herbivore density leads to less plant biomass in the plant-herbivore community, which in turn makes it more difficult for pollinators to invade (**Fig. S6e vs S6h, point A**). Finally, an increase in the herbivore conversion efficiency strengthens the related negative feedback loop, favoring stable dynamics (**Fig. S6a vs S6d, point C**).

#### 2. Analytical grounding

Animal conversion efficiencies affect both the feasibility and stability of coexistence (relationships (1) (2) (3) (4)). We determine here how relationships (1) (2) (3) vary with conversion efficiencies $e_m$ and $e_h$.

$$\frac{\partial(r_m + e_m a_{pm} P_{PH}^*)}{\partial e_m} = a_{pm} P_{PH}^* \geq 0$$

An increase in their conversion efficiency makes pollinators able to invade the plant-herbivore community more easily.

$$\frac{\partial(r_m + e_m a_{pm} P_{PH}^*)}{\partial e_h} = e_m a_{pm} \frac{\partial P_{PH}^*}{\partial e_h} = \frac{-e_m a_{pm} a_{ph}^2 P_{PH}^*}{c_p c_h + e_h a_{ph}^2} \leq 0$$

$$\frac{\partial H_{PH}^*}{\partial e_h} = \frac{a_{ph} r_p (c_p c_h + e_h a_{ph}^2) - (e_h a_{ph} r_p + c_p r_h) a_{ph}^2}{(c_p c_h + e_h a_{ph}^2)^2} = \frac{a_{ph} c_p (c_h r_p - a_{ph} r_h)}{(c_p c_h + e_h a_{ph}^2)^2}$$

$$= \frac{a_{ph} c_p P_{PH}^*}{c_p c_h + e_h a_{ph}^2} \geq 0$$

At the plant-herbivore equilibrium, the plant population gets smaller as a result of an increased herbivore density when herbivores are more efficient at exploiting their resource. It thus becomes harder for pollinators to invade the plant-herbivore community.

$$\frac{\partial\left[(c_p c_m - e_m {a_{pm}}^2)(r_h + e_h a_{ph} P_{PM}^*)\right]}{\partial e_m} = e_h a_{ph} \frac{\partial P_{PM}^*}{\partial e_m} = e_h a_{ph} a_{pm}^2 P_{PM}^* \geq 0$$

$$\frac{\partial M_{PM}^*}{\partial e_m} = \frac{c_p a_{pm} P_{PM}^*}{c_p c_m - e_m a_{pm}^2}$$

Relationship (2) is favored by an increase in the pollinator conversion efficiency (when $P_{PM}^* \leq 0$, relationship (2) is either always true or false (**Table S5**) so that we do not consider such cases here). This ensues from an increase in both plant and pollinator densities when the pollination intensity is too low to trigger unbounded growth (relationship (2'a)). It then becomes easier for herbivores to invade the sub-community. When pollination is strong enough for an orgy to occur in the plant-pollinator sub-community (relationship (2'b)), the relationship satisfaction is also favored by increasing $e_m$, although not mediated by the same positive effect on pollinator density.

$$\frac{\partial\left[(c_p c_m - e_m {a_{pm}}^2)(r_h + e_h a_{ph} P_{PM}^*)\right]}{\partial e_h} = (c_p c_m - e_m {a_{pm}}^2) a_{ph} P_{PM}^*$$

Relationship (2) is favored (resp. disfavored) by an increase in the herbivore conversion efficiency when unbounded PM growth is not possible (resp. possible) (see relationship (2'a) vs. (2'b)).

$$\frac{\partial(c_h c_m c_p + c_m e_h {a_{ph}}^2 - c_h e_m {a_{pm}}^2)}{\partial e_m} = -c_h {a_{pm}}^2 \leq 0$$

$$\frac{\partial(c_h c_m c_p + c_m e_h {a_{ph}}^2 - c_h e_m {a_{pm}}^2)}{\partial e_h} = c_m {a_{ph}}^2 \geq 0$$

Stability is favored when either the herbivore conversion efficiency increases or the pollinator conversion efficiency decreases.
